# DAZAP2 functions as a pan-coronavirus restriction factor by inhibiting viral entry and genomic replication

**DOI:** 10.1101/2025.02.04.636569

**Authors:** Fei Feng, Jiannan Chen, Rong Li, Yunkai Zhu, Yanlong Ma, Ziqiao Wang, Yuyan Wang, Zhichao Gao, Lulu Yang, Yin Yu, Yanfeng Liu, Yingjie Sun, Ying Liao, Xinxin Huang, Qisheng Zhang, Yongheng Huang, Lin Qiu, Jiayu Wu, Jingxian Zhao, Chao Liu, Qiang Ding, Youhua Xie, Zhenghong Yuan, Yue Hong, Ping Zhang, Jing Sun, Jincun Zhao, Rong Zhang

## Abstract

The SARS-CoV-2 pandemic and the emergence of novel variants underscore the need to understand host-virus interactions and identify host factors that restrict viral infection. Here, we perform a genome-wide CRISPR knockout screen to identify host restriction factors for SARS-CoV-2, revealing DAZAP2 as a potent antiviral gene. DAZAP2, previously implicated in SARS-CoV-2 restriction, is found to inhibit viral entry by blocking virion fusion with both endolysosomal and plasma membranes. Additionally, DAZAP2 suppresses genomic RNA replication without affecting the primary translation of viral replicases. We demonstrate that DAZAP2 functions as a pan-coronavirus restriction factor across four genera of coronaviruses. Importantly, knockout of *DAZAP2* enhances SARS-CoV-2 infection in mouse models and in human primary airway epithelial cells, confirming its physiological relevance. Mechanistically, antiviral activity of DAZAP2 appears to be indirect, potentially through the regulation of host gene expression, as it primarily localizes to the nucleus. Our findings provide new insights into the host defense system against coronaviruses and highlight DAZAP2 as a potential target for host-directed antiviral therapies.

**IMPORTANCE:** During viral infection, the host defense response is mediated by a variety of host factors through distinct mechanisms that have yet to be fully elucidated. Although *DAZAP2* was previously implicated in SARS-CoV-2 restriction, its mechanisms of action and *in vivo* relevance remain unclear. In this study, we identify the DAZAP2 as a potent pan-coronavirus restriction factor that inhibits viral infection through dual mechanisms: blocking virion fusion with both endolysosomal and plasma membranes, and suppressing genomic RNA replication. We confirm its physiological relevance in host defense using mouse models and primary cell cultures. This study advances our understanding of host-pathogen interactions. Targeting DAZAP2 or its regulatory pathways could provide a new approach to enhance host defense against current and future coronavirus threats.

## INTRODUCTION

*Coronaviridae* is a large family of enveloped, positive-sense, single-stranded RNA viruses and consists of four genera: *Alphacoronavirus*, *Betacoronavirus*, *Gammacoronavirus*, and *Deltacoronavirus.* Coronaviruses infect a variety of hosts, including humans, pigs, chickens, and other animals. One such virus, SARS-CoV-2, is the causative agent of the COVID-19 pandemic^1,2^. There are seven widely recognized human coronaviruses and although four of them (HCoV-229E, HCoV-NL63, HCoV-OC43, and HCoV-HKU1) cause the common cold, three (MERS-CoV, SARS-CoV, and SARS-CoV-2) emerged as major public health concerns in this century by causing severe infection with high morbidity and mortality^3–5^. In addition, some animal coronaviruses, such as porcine epidemic diarrhea virus (PEDV), swine acute diarrhea syndrome coronavirus (SADS-CoV), porcine deltacoronavirus (PDCoV), and infectious bronchitis virus (IBV), have led to significant economic loss in the agricultural industry^6^. Closely related to a bat coronavirus, SADS-CoV was first identified in pigs and can infect primary human lung and intestinal cells^7^. Furthermore, infections with PDCoV have been detected among young children in Haiti^8^. Thus, coronaviruses pose a broad and considerable threat to public health, social stability, and the economy.

Coronaviruses have large RNA genomes of 28-31 kb, encoding for multiple structural and nonstructural proteins that have conserved functions in the infection cycles of different coronaviruses^4^. As a result, coronaviruses usually employ and share the same host machinery to facilitate infection and, conversely, can be met with similar host defense responses to restrict the infection. A great effort has been made by scientists to understand the anti-coronavirus host response, particularly to identify restriction factors of SARS-CoV-2. Interferons (IFNs) play a key role during virus infection, and IFN-stimulated genes (ISGs) often possess antiviral activities^9,10^. ISGs, such as LY6E, BST2, DAXX, OAS1, and many others against SARS-CoV-2 infection, have been identified either by arrayed ISG cDNA overexpression^11–13^, or by focused CRISPR knockout or activation library of ISGs^14,15^. In addition to ISGs, host restriction factors like mucins, are identified for SARS-CoV-2 by genome-scale CRISPR activation strategy^16,17^. MHC class II transactivator CIITA-induced cell resistance to SARS-CoV-2 infection has been reported by using a transposon-mediated gene-activation screen^18^. Moreover, genome-wide association studies as well as transcriptomic and proteomic analyses of SARS-CoV-2-infected cell lines and patient samples have uncovered additional host factors that may restrict or confer resistance to viral infection^19–28^.

To comprehensively identify host restriction factors of SARS-CoV-2, we conducted a genome-wide CRISPR/Cas9 knockout screen coupled with fluorescence-activated cell sorting (FACS) to enrich for susceptible cells. This approach revealed a suite of host factors with antiviral activity, including DAZ-associated protein 2 (DAZAP2), which was previously identified as an antiviral factor in a CRISPR dropout screen by analyzing the depleted cells^29^. Although DAZAP2 was shown to restrict SARS-CoV-2 infection, potentially through regulation of SERPINE1 expression^29^, how it impacts the life cycle of virus infection and whether it functions *in vivo* remain to be elucidated. In this study, we demonstrate that DAZAP2 is a pan-coronavirus restriction factor that acts at the stages of viral entry and replication. Importantly, we validated its antiviral activity in mouse models and human primary airway epithelial cells, where knockout of DAZAP2 significantly enhanced SARS-CoV-2 infection. These findings provide insights into the host defense mechanisms against coronaviruses and highlight DAZAP2 as a potential target for broad-spectrum antiviral strategies.

## RESULTS

### DAZAP2 is a pan-coronavirus host restriction factor

To identify host factors that can restrict SARS-CoV-2 infection, we performed a genome-wide, cell sorting-based screen in ACE2-expressing A549 (A549-ACE2) transduced with a knockout library of single-guide RNAs (sgRNAs) targeting 19,114 human genes^30^. The rationale is that knockout of antiviral host genes will enhance the virus infection. The library-transduced cells were infected with transcription- and replication-competent SARS-CoV-2 virus-like particles where the N protein was replaced with GFP (SARS-CoV-2 trVLP-GFP), resulting in only single-cycle infection^31^. Unlike previous studies that analyzed depleted cells to identify antiviral genes ^29^, we sorted virus-infected GFP-positive cells for genomic DNA extraction, sgRNA sequencing, and data analysis (Supplementary table 1). The genes identified from the screen were ranked based on the MAGeCK score (Fig. 1A). The top hit was programmed cell death 10 (*PDCD10* or *CCM3*), a gene associated with apoptosis^32,33^, followed by *DAZAP2*, which encodes a multifunctional proline-rich protein involved in cell signaling^34,35^, transcription regulation^36,37^, stress granule formation^38^. Other highly ranked genes included calcium-binding protein 39 (*CAB39* or *MO25*), phospholipid scramblase 1 (*PLSCR1*), vesicle trafficking 1 (*VTA1* or *LIP5*), and Lymphocyte antigen 6E (*LY6E*). *DAZAP2* and *VTA1* were previously identified as SARS-CoV-2 restriction factors in a CRISPR dropout screen, potentially through regulation of SERPINE1 expression ^29^. *PLSCR1* is an antiviral gene strongly induced by virus infection or interferon treatment^39–41^, and was previously identified as a host factor associated with critical COVID-19^27^. *PLSCR1* is also reported to restrict the SARS-CoV-2 infection at a late entry step before viral RNA is released into host cells^42^. *CAB39*, a component of a trimeric complex including serine/threonine kinase 11 (*STK11*) and STE20-related adaptor alpha (*STRAD*), has been studied in the context of cancer progression^43,44^, but with unknown function during SARS-CoV-2 infection. *VTA1* is involved in trafficking of endosomal multivesicular bodies and retrovirus budding^45^.

**Fig. 1.**
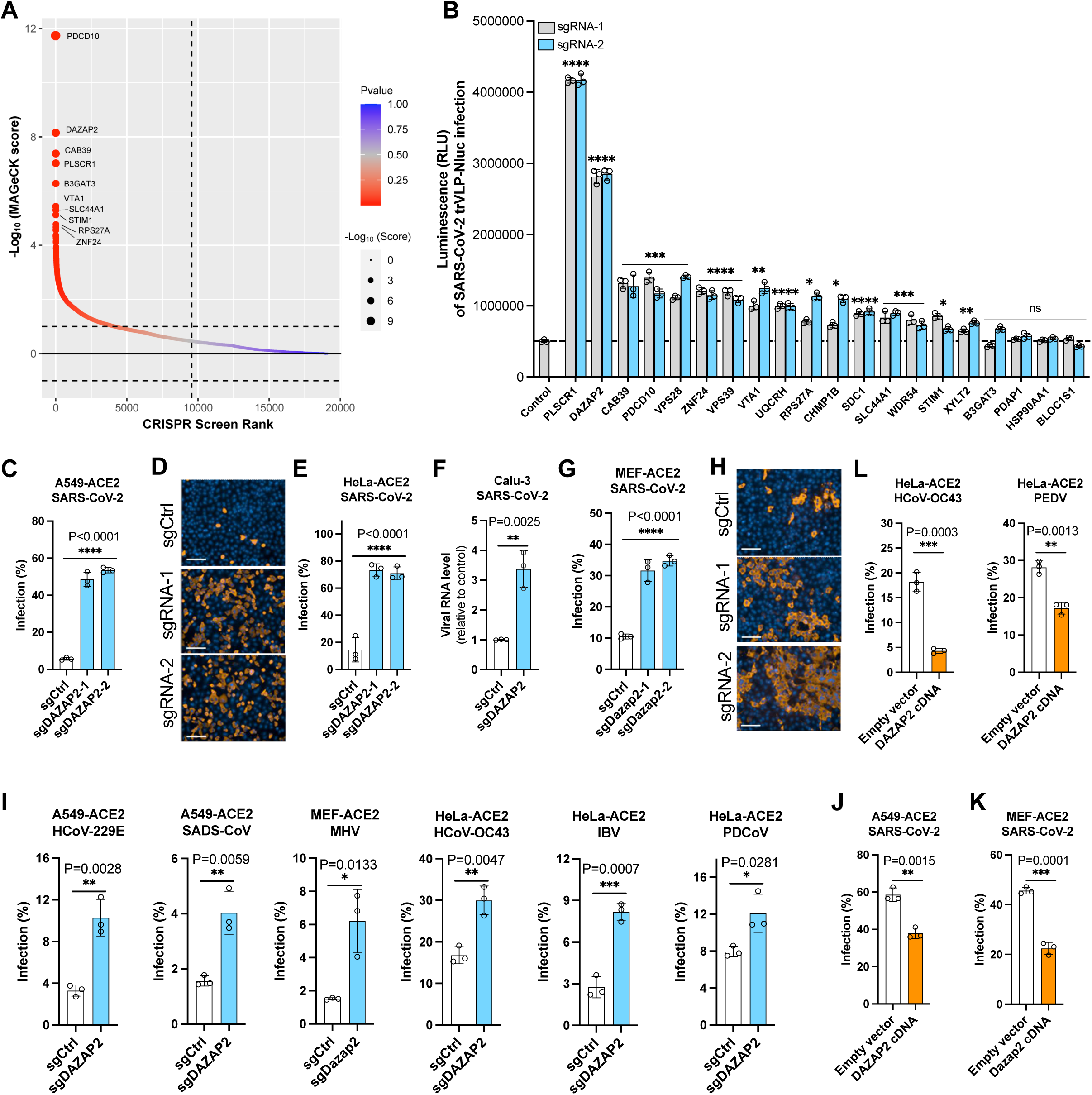
DAZAP2 is a pan-coronavirus host restriction factor. **A.** Genes identified from the CRISPR screen. ACE2-expressing A549 (A549-ACE2) cells transduced with a CRISPR knockout library were infected with SARS-CoV-2 transcription-and replication-competent virus-like particles where the N gene is replaced by the reporter GFP (trVLP-GFP) (MOI 0.5, 24 h). trVLP-GFP was packaged in cells expressing the N gene and only replicated for single round in A549-ACE2 in the absence of N protein. GFP-positive cells were sorted for genomic extraction and sgRNA sequence analysis. The genes were analyzed by MAGeCK software and sorted based on the -log_10_ (MAGeCK score). **B.** Experimental validation of top 20 genes from the screen in A549-ACE2 cells. Two independent sgRNAs per gene were used, and cells were infected with SARS-CoV-2 transcription- and replication-competent virus-like particles where the N gene is replaced by the NanoLuc luciferase (trVLP-Nluc) (MOI 0.5, 24 h). The infection efficiency was quantified by measuring the luciferase activity. **C.** High content imaging and quantification analysis of SARS-CoV-2 infection in *DAZAP2*-edited A549-ACE2 (MOI 0.1, 24 h). **D.** Representative immunofluorescence images of SARS-CoV-2 infection in *DAZAP2*-edited A549-ACE2 cells (MOI 0.1, 24 h). Scale bar, 100 μm. **E.** High content imaging and quantification analysis of SARS-CoV-2 infection in *DAZAP2*-edited HeLa-ACE2 (MOI 0.1, 24 h). **F.** qRT-PCR was conducted to measure the N gene copies of SARS-CoV-2 infected Calu-3 cells (MOI 1, 24 h). GAPDH was used as internal control. **G-H.** High content imaging and quantification analysis (**G**) and immunofluorescence images (**H**) of SARS-CoV-2 infection in mouse *Dazap2*-edited MEFs expressing human ACE2 (MEF-ACE2) (MOI 0.1, 24 h). Scale bar, 100 μm. **I.** Validation of DAZAP2 as a restriction factor during infection with other coronaviruses. Gene-edited cells were infected with alphacoronaviruses (HCoV-229E, MOI 1, 48 h; SADS-CoV, MOI 3, 24 h), betacoronaviruses (MHV, MOI 5, 24 h; HCoV-OC43, MOI 0.03, 12 h), gammacoronaviruses (IBV, MOI 0.5, 24 h), or deltacoronaviruses (PDCoV, MOI 0.3, 24 h). **J-K.** Overexpression of DAZAP2 inhibits SARS-CoV-2 infection (MOI 1, 24 h). Human *DAZAP2* or mouse *Dazap2* cDNA was expressed in A549-ACE2 or MEF-ACE2, respectively. **L.** Overexpression of human DAZAP2 in HeLa-ACE2 inhibits HCoV-OC43 (MOI 0.03, 24 h) and PEDV (MOI 1, 24 h) infection. The virus infection efficiency was determined by analyzing the percentage of viral N-positive cells using flow cytometry or Operetta High Content Imaging System. Data shown are from three independent experiments and each independent experiment was performed in duplicate or triplicate. B, two-way ANOVA with Dunnett’s test; the mean of two sgRNAs was compared with the control sgRNA; C, E, and G, one-way ANOVA with Dunnett’s test; F and I-L, unpaired t test; n=3; mean ± s.d.; *P < 0.05; **P < 0.01; ***, P < 0.001; ****P < 0.0001; ns, not significant.

To validate the candidate hits, we chose the 20 top-ranked genes. For each specific gene target, A549-ACE2 cells were edited with two independent sgRNAs. We also modified the transcription-and replication-competent SARS-CoV-2 virus-like particle (SARS-CoV-2 trVLP) where the N gene was replaced by NanoLuc luciferase instead of GFP (SARS-CoV-2 trVLP-Nluc) (Supplementary Fig. 1). Thus, infection efficiency could be assessed by quantifying luciferase activity over time. Knockout of the antiviral gene *PLSCR1* had the greatest effect, resulting in an 8-fold increase of infectivity when compared to control (Fig. 1B). *DAZAP2* knockout increased infectivity by over 4 folds. Knockout of *CAB39*, *PDCD10*, *VPS28*, *VPS39*, *VTA1*, *ZNF28*, or *UQCRH* each resulted in up to 2-fold increases in infectivity compared to control.

Of the validated genes, *DAZAP2* strongly restricted SARS-CoV-2 infection, consistent with previous findings^29^. However, how DAZAP2 impacts the life cycle of coronavirus infection remains to be elucidated. We therefore focused the rest of our study on *DAZAP2,* attempting to understand its role in coronavirus infection. We first confirmed its antiviral activity of *DAZAP2* in the context of infection with authentic SARS-CoV-2. Two independent sgRNAs were used to knockout *DAZAP2* in A549-ACE2 cells. The infection efficiency was increased by approximately 10 folds, and the representative images of virus infection were indicated (Fig. 1C and D). Similarly, SARS-CoV-2 infection in ACE2-expressing HeLa cells (HeLa-ACE) was increased by approximately 5 folds when the DAZAP2 was edited (Fig. 1E). We further validated the role of *DAZAP2* as a restriction factor in Calu-3 cells, a physiologically relevant lung epithelial cell line, by quantifying viral RNA levels in cells (Fig. 1F). Additionally, knockout of mouse orthologue *Dazap2* in murine embryonic fibroblasts expressing human ACE2 (MEF-ACE2) significantly enhanced SARS-CoV-2 infection (Fig. 1G and H). Knockout efficiency of *DAZAP2* was confirmed by western blotting in all cell lines (Supplementary Fig. 2A).

To determine whether *DAZAP2* acts as a pan-coronavirus restriction factor, we infected *DAZAP2*-edited A549-ACE, HeLa-ACE2, and MEF-ACE2 with viruses from all four *Coronaviridae* genera: the alphacoronaviruses HCoV-229E and SADS-CoV; the betacoronaviruses HCoV-OC43 and mouse hepatitis virus (MHV); the gammacoronavirus IBV; and the deltacoronavirus PDCoV. *DAZAP2* knockout significantly enhanced infection by all tested coronaviruses (Fig. 1I). Conversely, overexpression of either human or mouse *DAZAP2* suppressed SARS-CoV-2 infection (Fig. 1J and K). Protein expression of exogenous human DAZAP2 cDNA was verified by western blotting (Supplementary Fig. 2B). Similarly, overexpression of human *DAZAP2* inhibited infection by the betacoronavirus HCoV-OC43 and the alphacoronavirus porcine epidemic diarrhea virus (PEDV) (Fig. 1L). Together, these data demonstrate that DAZAP2 is a conserved, pan-coronavirus restriction factor that functions across cell types and species.

### DAZAP2 inhibits the endosomal entry of SARS-CoV-2

Having established DAZAP2 as a common restriction factor for coronaviruses across four genera, we next sought to define its role in the coronavirus life cycle. To determine the stage of infection at which DAZAP2 acts, we infected *DAZAP2*-edited cells with murine leukemia retrovirus (MLV)-based pseudoviruses bearing the spike protein of SARS-CoV-2 or, as a control, the glycoprotein of vesicular stomatitis virus (VSV-G). Knockout of *DAZAP2* resulted in an approximately 25-fold increase in SARS-CoV-2 pseudovirus infection, while no significant difference was observed for the VSV-G pseudovirus (Fig. 2A and B). Similarly, pseudoviruses packaged with the spike from SARS-CoV-1 exhibited about 9-fold increase in infection, consistent with the results for SARS-CoV-2 pseudoviruses (Fig. 2C).

**Fig. 2.**
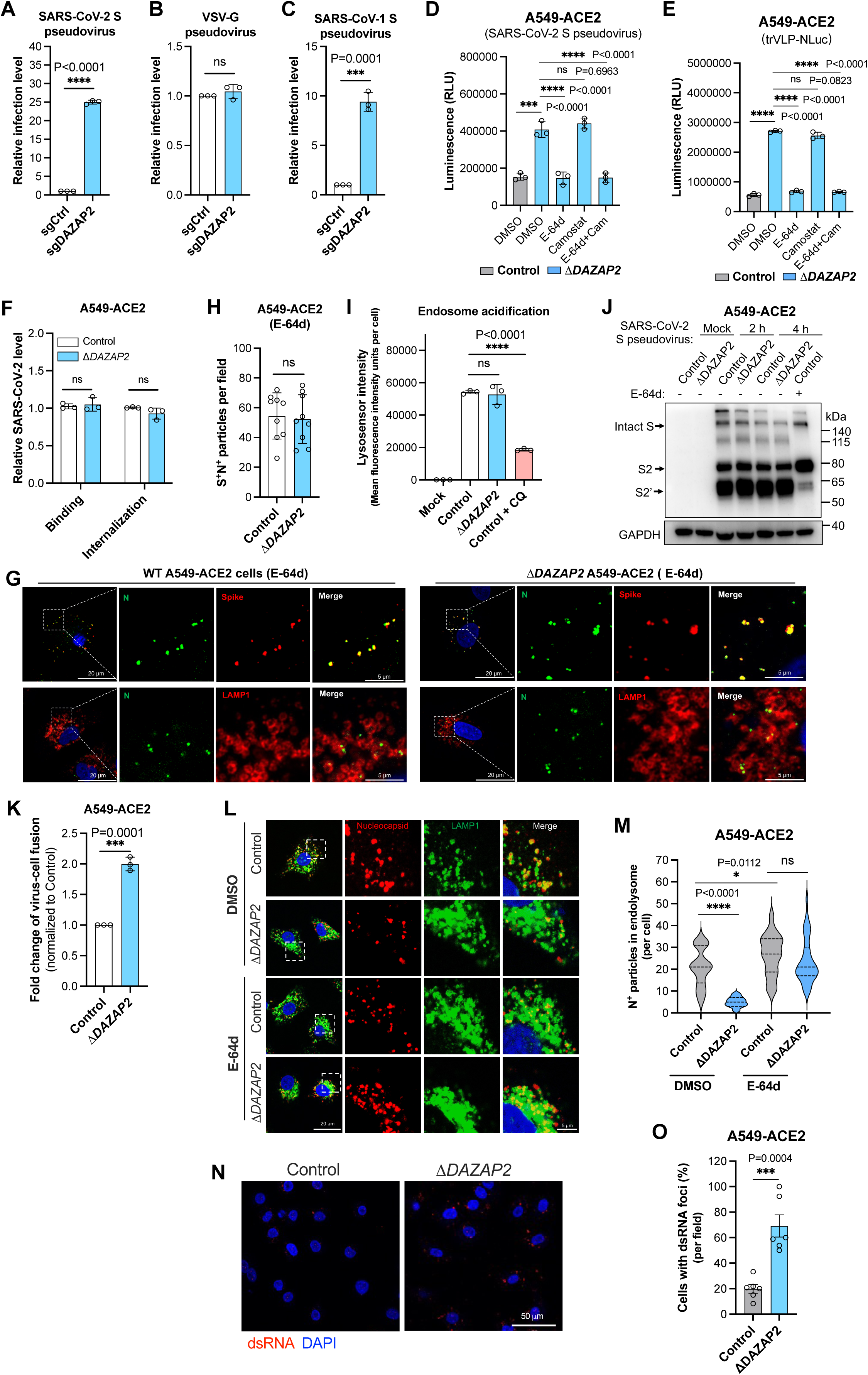
DAZAP2 inhibits virion fusion with endolysosome membranes to release genomes into the cytosol. **A-C.** Pseudovirus infection assay. The gene-edited A549-ACE2 cells were infected with murine leukemia retrovirus (MLV)-based pseudoviruses bearing the spike protein of SARS-CoV-2 (**A**), the glycoprotein of vesicular stomatitis virus (VSV-G) (**B**), or the spike protein of SARS-CoV-1 (**C**), and the luciferase activity was measured and normalized to the control. **D-E.** The inhibition of endosomal entry of SARS-CoV-2. Control and *DAZAP2*-knockout clonal cell line of A549-ACE2 (11*DAZAP2*) were infected with MLV-based pseudovirus bearing the spike protein of SARS-CoV-2 (D) or single-cycle trVLP-NLuc (E), in the presence of 100 μM E-64d (aloxistatin), an inhibitor that blocks the cysteine protease activity of cathepsins B and L, which are required for the endosomal membrane fusion, and/or 100 μM camostat mesylate, a TMPRSS2 inhibitor that blocks viral fusion at the plasma membrane. **F.** Virus binding and internalization assays. Cells were incubated with SARS-CoV-2 (MOI 5), and the bound or internalized virions were measured by qRT-PCR for genomic RNA. **G-H.** Trafficking of SARS-CoV-2 trVLP-Nluc particles in the presence of cysteine protease inhibitor E-64d (100 μM). The representative confocal images (G) were obtained, and the quantification of spike and N protein double-positive particles per field (H) was analyzed. The endolysosome marker LAMP1 was stained. Scale bar, 20 or 5 μm. **I.** Quantification of endosomal acidification. Control or *DAZAP2*-deficient A549-ACE2 cells were pre-treated with or without 20 μM chloroquine (CQ) followed by staining of LysoSensor Green dye. The fluorescence intensity was quantified. **J.** The cleavage of the SARS-CoV-2 spike protein. Control or *DAZAP2*-deficient A549-ACE2 cells were incubated with MLV-based pseudoviruses bearing the spike protein for 2 or 4 h, followed by western blotting analysis with anti-S2 antibody. The cysteine protease inhibitor E-64d (100 μM) was used as control. **K.** Split NanoLuc luciferase reporter-based virus-cell fusion assay. Cells expressing the LgBit were incubated with retrovirus particles encapsulated with CypA-HiBit to enable virion fusion in the endolysosomes. The re-complemented NanoLuc luciferase activity in the cytoplasm was determined and normalized to the control. **L-M.** Quantification of virions in the endolysosomes. Control or *DAZAP2*-deficient A549-ACE2 cells were infected with SARS-CoV-2 for 4 h, and fixed to stain the spike, N, and endolysosome marker LAMP1. The colocalization of LAMP1 with N was visualized by confocal microscopy (L) and the number of colocalized foci per cell was counted (M). Three fields of view with a total of 27 to 42 cells were used for analysis. The representative confocal images (L) were shown. Scale bar, 20 or 5 μm. The cysteine protease inhibitor E-64d (100 μM) was used as control. **N-O.** Quantification of double-stranded RNA (dsRNA). Control or *DAZAP2*-deficient A549-ACE2 cells were infected with SARS-CoV-2 for 4 h, then fixed to stain the dsRNA with J2 antibody. The dsRNA puncta were visualized by confocal microscopy (N) and the percentage of dsRNA-positive cells per field was counted (O). Six fields of view with a total of 109 control cells and 79 *DAZAP2*-deficient cells were selected for analysis. The representative confocal images (N) were shown. Scale bar, 50 μm. Data shown are from three independent experiments and each independent experiment was performed in triplicate. A-C, F, H, K, and O, unpaired t test; D, E, I, and M, one-way ANOVA with Dunnett’s test; n=3; mean ± s.d.; *P < 0.05; ***, P < 0.001; ****P < 0.0001; ns, not significant.

Since A549 cells express little or no TMPRSS2, a serine protease that facilitates viral entry via plasma membrane fusion, SARS-CoV-2 primarily enters A549-ACE2 cells through the endosomal pathway. To test whether DAZAP2 inhibits endosomal entry, we generated and validated a *DAZAP2*-knockout clonal cell line (Δ*DAZAP2*) of A549-ACE2 (Supplementary Fig. 2C to E). Control or Δ*DAZAP2* cells were infected with SARS-CoV-2 pseudovirus in the presence of E-64d (aloxistatin), an inhibitor of the cysteine proteases cathepsins B and L (required for endosomal membrane fusion), and/or camostat mesylate, a TMPRSS2 inhibitor that blocks plasma membrane fusion (Fig. 2D). We found that the enhancement of pseudovirus infection in Δ*DAZAP2* cells is significantly diminished in the presence of E-64d, whereas camostat mesylate had no effect. Similar results were observed when cells were infected with single-cycle SARS-CoV-2 trVLP-Nluc particles. These findings suggest that DAZAP2 primarily inhibits the endosomal entry pathway of SARS-CoV-2.

### DAZAP2 inhibits virion fusion with endolysosome membranes to release genomes into the cytosol

To further dissect the SARS-CoV-2 entry process, we divided viral entry into four distinct steps: (1) binding, (2) uptake (internalization), (3) virion trafficking to late endosome/lysosomes, and (4) virion fusion with the endolysosome membranes to release viral genomes into the cytosol. A previous study indicated that DAZAP2 does not affect the binding and internalization of SARS-CoV-2^29^, which initially appeared contradictory to our results (Fig. 2A to E). To resolve this, we incubated virions on ice for 45 min with control and Δ*DAZAP2* cells to assess binding. For internalization, cells were shifted to 37°C for 45 min after the binding to allow virion uptake. Consistent with prior findings, neither binding to the cell surface nor internalization was affected in *DAZAP2*-deficient cells (Fig. 2F).

Next, we investigated whether DAZAP2 influences the trafficking of internalized virions to endolysosomes. After internalization at 37°C for 4 hours in the presence of E-64d (to prevent virion fusion with endolysosome membranes), we quantified virions (spike-positive and nucleocapsid-positive) co-localized with the endolysosome marker LAMP1. No significant difference in virion counts was observed between control and 11*DAZAP2* cells (Fig. 2G and H).

We then examined whether DAZAP2 affects virion fusion with endolysosomes, a process requiring the activation of cysteine proteases (cathepsins B and L) in a low pH environment. Using LysoSensor Green dye, we found that endolysosomal acidification was unchanged in Δ*DAZAP2* cells (Fig. 2I). Similarly, the cleavage of the spike protein by cysteine proteases was unaffected in Δ*DAZAP2* cells (Fig. 2J). These results demonstrate that DAZAP2 does not impact SARS-CoV-2 endosomal entry at the stages of binding, uptake, trafficking, or spike cleavage.

The final step in endosomal entry is the fusion of virions with endolysosome membranes to release viral genomes into the cytosol. To assess this, we employed an improved virus-cell fusion assay^42^. Cyclophilin A (CypA), a Gag-interacting protein, was fused with HiBit and encapsulated into retrovirus particles bearing the SARS-CoV-2 spike protein. If internalized virions fuse with endolysosome membranes, the released HiBit into the cytosol complements LgBit to form functional NanoLuc luciferase. When 11*DAZAP2* cells expressing LgBit were incubated with these particles, luciferase activity was significantly higher compared to control cells (Fig. 2K), indicating enhanced fusion in 11*DAZAP2* cells.

To further validate these findings, control and Δ*DAZAP2* cells were infected with authentic SARS-CoV-2 for 4 hours, followed by detection of virion particles co-localized with endolysosome marker LAMP1 (Fig. 2L). As expected, fewer virions were observed in 11*DAZAP2* cells when compared to controls (Fig. 2L and M). Conversely, intracellular double-stranded RNA (dsRNA), an indicator of viral replication intermediates, was more abundant in Δ*DAZAP2* cells (Fig. 2N and O). As a control, the addition of E-64d resulted in similar numbers of virions in endolysosomes for both cell types (Fig. 2L and M). These findings collectively suggest that DAZAP2 inhibits the fusion of SARS-CoV-2 virions with endolysosome membranes, thereby preventing the release of viral genomes into the cytoplasm.

### DAZAP2 restricts the plasma membrane entry of SARS-CoV-2

We next investigated whether DAZAP2 affects spike-protein-mediated fusion at the plasma membrane. Control and Δ*DAZAP2* A549-ACE2 acceptor cells were co-cultured with 293T donor cells expressing the SARS-CoV-2 spike protein, and spike-mediated cell-cell syncytia formation was visualized. Notably, Δ*DAZAP2* cells exhibited a larger fusion area and more syncytial nuclei compared to control cells (Fig. 3A and B).

**Fig. 3.**
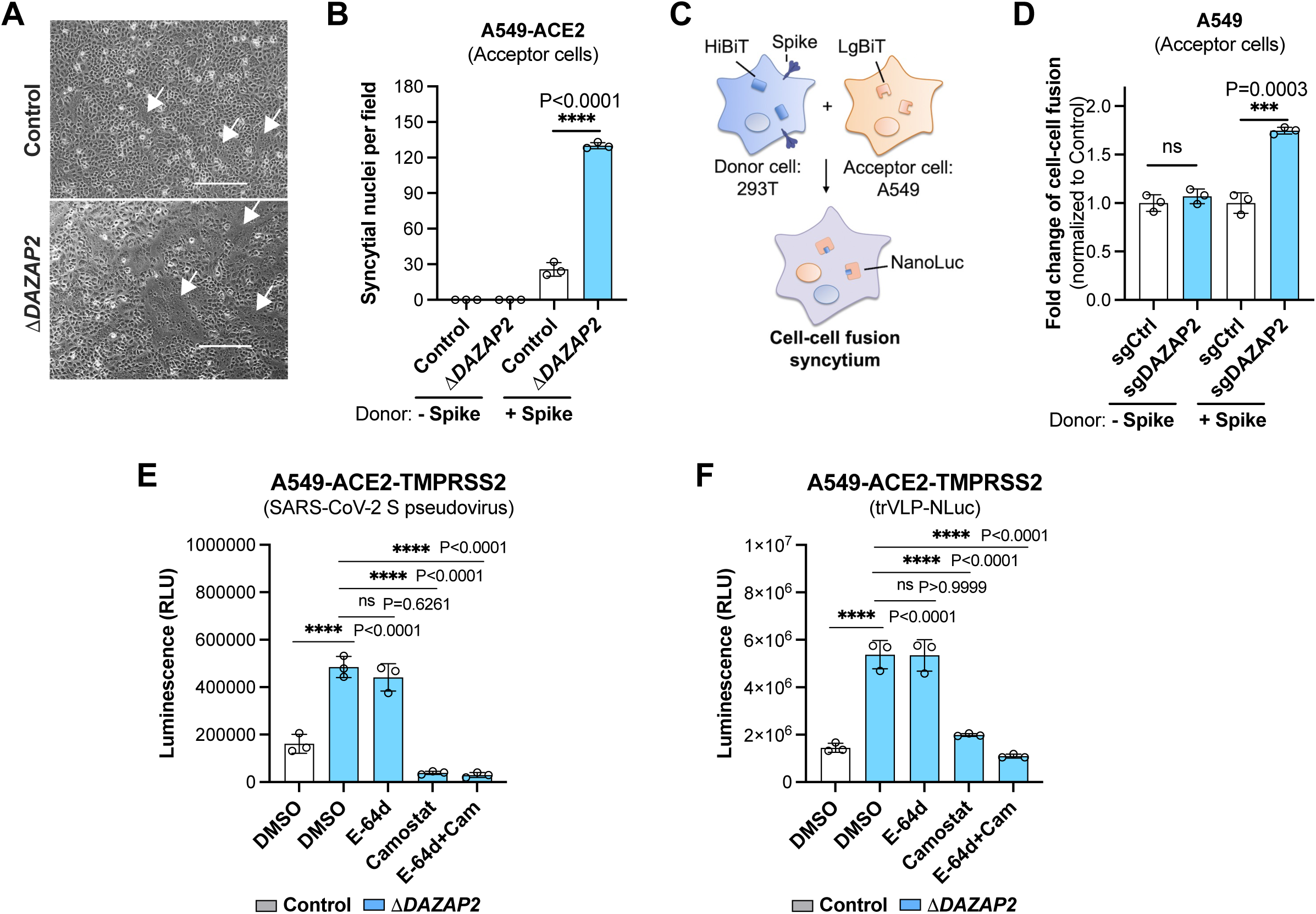
DAZAP2 inhibits the plasma membrane entry of SARS-CoV-2. **A-B.** Cell-cell fusion assay. Control and Δ*DAZAP2* A549-ACE2 acceptor cells were co-cultured with 293T donor cells that express SARS-CoV-2 spike protein. Spike protein-induced syncytia was visualized under brightfield microscope (A), and syncytial nuclei were counted after Giemsa staining (B). Scale bar, 100 μm. **C-D.** Schematic (C) and the results (D) of split NanoLuc luciferase reporter-based cell-cell fusion assay. A549-ACE2 acceptor cells expressing the LgBit were incubated with 293T donor cells expressing both HiBit and SARS-CoV-2 spike protein. The functional NanoLuc luciferase was re-complemented after cell-cell fusion and the activity was measured and normalized to the control. **E-F.** The inhibition of plasma membrane entry of SARS-CoV-2. Control and Δ*DAZAP2* A549-ACE2 cells ectopically expressing the TMPRSS2 (A549-ACE2-TMPRSS2) were infected with MLV-based pseudovirus bearing the spike protein of SARS-CoV-2 (E) or single-cycle trVLP-NLuc (F), in the presence of cysteine protease inhibitor E-64d (100 μM), and/or TMPRSS2 inhibitor camostat mesylate (100 μM). Data shown are from three independent experiments and each independent experiment was performed in triplicate. B and C, unpaired t test; E and F, one-way ANOVA with Dunnett’s test; n=3; mean ± s.d.; ***, P < 0.001; ****P < 0.0001; ns, not significant.

To quantify the cell-cell fusion, we employed a split NanoLuc luciferase-based assay as previously described^42^. In this system, acceptor cells express the LgBit fragment, while the donor cells express the HiBit fragment. Upon co-culture and spike-mediated cell-cell fusion, the two fragments complement each other to form functional NanoLuc luciferase, allowing for quantification of fusion activity (Fig. 3C). Consistent with the syncytia formation results (Fig. 3A and B), we observed a significant increase in luciferase activity in Δ*DAZAP2* cells, indicating enhanced cell-cell fusion (Fig. 3D).

Because A549 cells express minimal or no TMPRSS2 protease that is required for spike-mediated plasma membrane entry, we generated A549-ACE2-TMPRSS2 cells (control and Δ*DAZAP2*) to further explore this pathway. These cells were infected with SARS-CoV-2 pseudovirus in the presence of E-64d (to block endosomal entry) and/or camostat mesylate (a TMPRSS2 inhibitor) (Fig. 2D). In TMPRSS2-expressing cells, E-64d failed to inhibit infection, whereas the camostat mesylate significantly reduced the enhanced infection observed in Δ*DAZAP2* cells (Fig. 3E). Similar results were obtained using single-cycle SARS-CoV-2 trVLP-Nluc particles (Fig. 3F). These results demonstrate that DAZAP2 inhibits the SARS-CoV-2 spike protein-mediated cell-cell fusion and plasma membrane entry, further underscoring its role as a broad-spectrum restriction factor.

### DAZAP2 inhibits the genomic replication of SARS-CoV-2

The SARS-CoV-2 trVLP particles, which lack the N gene, recapitulate only the viral entry and replication stages of the life cycle, excluding virion assembly and release. Consequently, the CRISPR screen using SARS-CoV-2 trVLP is biased toward identifying host factors involved in entry and replication. In addition to its role in inhibiting viral entry we sought to determine whether DAZAP2 also affects post-entry steps, particularly viral genomic replication.

To investigate this, we constructed a SARS-CoV-2 replicon system in which the portion of the genome encoding the spike protein all the way through ORF8 was replaced by NanoLuc luciferase (Supplementary Fig.1). The *in vitro* transcribed replicon RNA was electroporated into cells, and viral replication was monitored by measuring the luciferase activity. Knockout of *DAZAP2* led to significantly enhanced viral replication when compared to control cells (Fig. 4A). Treatment with remdesivir, an RNA-dependent RNA polymerase (RdRp) inhibitor, diminished the luciferase activity in both control and *DAZAP2*-edited cells, confirming the utility of this system to assess changes in replication (Fig. 4A). Conversely, overexpression of DAZAP2 could significantly inhibit the replication (Fig. 4B).

**Fig. 4.**
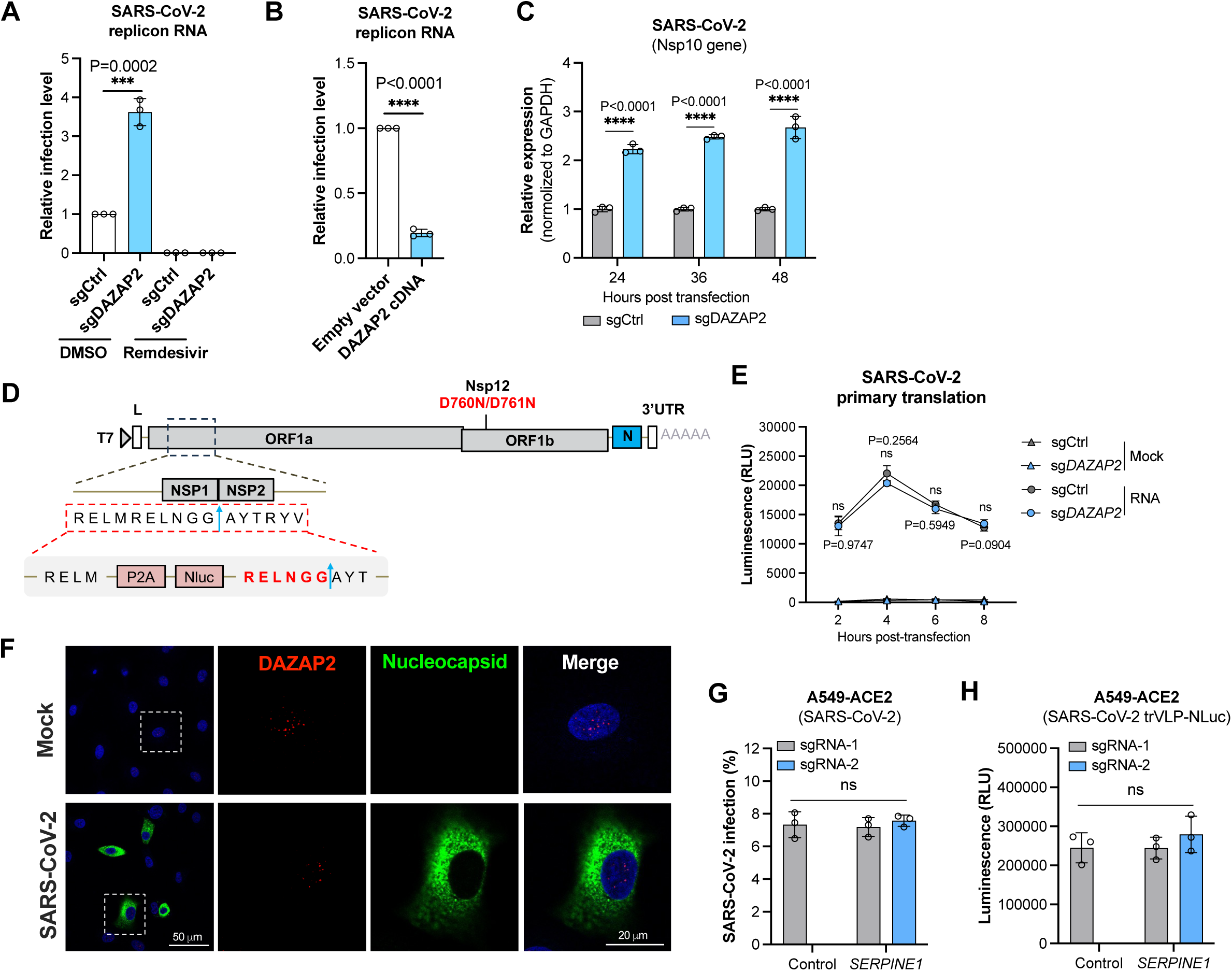
DAZAP2 inhibits the genomic replication of SARS-CoV-2. **A.** Replicon RNA assay in HeLa cells edited with control or *DAZAP2* sgRNA. The SARS-CoV-2 replicon system was constructed by replacing the portion of the genome encoding the spike protein all the way through ORF8 with NanoLuc luciferase. The *in vitro* transcribed replicon RNA was electroporated into cells. The RNA-dependent RNA polymerase (RdRp) inhibitor remdesivir (10 μM) was added as a control to verify the utility of the replicon system. One representative sgRNA was used to edit the *DAZAP2*. The luciferase activity was determined and normalized to the control. **B.** Replicon RNA assay in empty vector-and *DAZAP2*-overexpressing HeLa cells, and the results were normalized to the control. **C.** Quantification of genomic RNA replication. The *in vitro* transcribed replicon RNA was electroporated into cells, and the levels of genomic RNA replication was determined by qRT-PCR targeting the NSP10 gene at the indicated timepoints. The results were normalized to the control. **D.** Schematic of the construction of inactivated replicon system to assess the primary translation of viral replicases. The NanoLuc luciferase reporter gene, flanked by a P2A cleavage site, was inserted between NSP1 and NSP2 of the SARS-CoV-2 replicon. The D760N and D761N double mutations were introduced into the NSP12 to inactivate the RdRp activity, ensuring that only translation could be assessed. **E.** Detection of the primary translation of viral replicases as indicated by the luciferase activity. Control and DAZAP2-edited HeLa cells were electroporated with the modified replicon RNA, and the luciferase activity was monitored. **F.** Confocal analysis of the localization of DAZAP2 and SARS-CoV-2 N protein. SARS-CoV-2-infected A549-ACE2 cells were fixed and stained with anti-DAZAP2 or anti-N antibody. The representative confocal images were shown. Scale bar, 50 or 20 μm. **G-H.** Validation of *SERPINE1* gene. A549-ACE2 cells were edited with two independent sgRNAs targeting the *SERPINE1*, followed by infection with authentic SARS-CoV-2 (G) or trVLP-NLuc particles (H). Data shown are from three independent experiments and each independent experiment was performed in triplicate. As for the results shown as relative change, data are normalized to the control of individual experiment. A-C, G and H, unpaired t test; E, two-way ANOVA; n=3; mean ± s.d.; ****P < 0.0001; ns, not significant.

To further validate these findings, we quantified genomic RNA levels by qRT-PCR targeting the NSP10 gene at different timepoints. Higher levels of genomic RNA were detected in *DAZAP2*-edited cells than that in control (Fig. 4C). These results suggest that DAZAP2 inhibits viral genomic RNA replication, leading to the reduced protein translation, as indicted by decreased NanoLuc luciferase activity.

We next explored whether the inhibition of genomic replication by DAZAP2 is due to its impact on the primary translation of non-structural replicases for incoming viral genomes released from endolysosomes. To test this, we modified the replicon system by inserting the NanoLuc luciferase reporter gene, flanked by a P2A cleavage site, between NSP1 and NSP2 of the SARS-CoV-2 replicon (Fig. 4D). Additionally, we introduced D760N and D761N double mutations into the NSP12 to inactivate the RdRp activity^46^, ensuring that only translation could be assessed (Fig. 4D). After electroporating the modified replicon RNA into cells, the luciferase activity was monitored from 2 to 8 hours, peaking at 4 hours (Fig. 4E). However, no significant difference in luciferase activity was observed between control and *DAZAP2*-edited cells, indicating that DAZAP2 does not affect the primary translation of viral replicases.

### DAZAP2 may regulate host gene expression to restrict viral infection

To understand how DAZAP2 inhibits coronavirus entry and replication, we first evaluated whether it is an interferon-stimulated gene (ISG). *DAZAP2* gene expression was not upregulated upon treatment with 1,000 U/ml IFNα-2b (Supplementary Fig. 3A) whereas the known ISGs IFITM3 and MX1 were increased over 100- and 1000-fold, respectively (Supplementary Fig. 3B and C). Furthermore, we performed SARS-CoV-2 infection in *STAT1*-, *MAVS*-, or *IRF3*-deficient A549-ACE2 cells, which lack critical innate immune signaling pathways. Despite these disruptions, infection efficiency was still significantly increased in *DAZAP2*-edited cells as compared to the controls (Supplementary Fig. 4). These results suggest that the antiviral function of DAZAP2 is possibly independent of innate immune responses.

We next investigated whether DAZAP2 directly exerts its antiviral function at the endolysosome or cytoplasm to inhibit viral entry and replication, respectively. Localization studies revealed that endogenous DAZAP2 is predominantly located in the nucleus in both mock and virus-infected cells, while viral N protein localizes to the cytoplasm (Fig. 4F). Additionally, DAZAP2 did not co-localize with the early endosome marker EEA1 or the endolysosome marker LAMP1 (Supplementary Fig. 5), further supporting that DAZAP2 may not directly interact with viral entry machinery at these sites.

Given that DAZAP2 localizes to the nucleus and has been reported to interact with transcription factors^36,37^, we hypothesized that it may regulate the expression of specific host genes, contributing to its role as a restriction factor. While Hou et al. previously suggested that DAZAP2, along with VTA1 and KFL5, regulates host cell responses to SARS-CoV-2 infection by controlling the *SERPINE1* expression (associated with COVID-19 severity), we found that editing *SERPINE1* had no effect on infection by authentic SARS-CoV-2 or trVLP-NLuc particles (Fig. 4G and H).

Since DAZAP2 inhibits viral entry at both the endosomal and plasma membranes, we also examined whether DAZAP2 affects the expression of SASR-CoV-2 entry factors. Using a validated antibody against the cell receptor ACE2 (Supplementary Fig. 6A and B), the surface expression level of ACE2 was not changed when the DAZAP2 was edited (Supplementary Fig. 6C). Likewise, the expression of other known entry factors, such as AXL, heparan sulfate, TIM-1, SIGLEC1, DC-SIGN, CTSL, Furin, and TMPRSS2, were not affected by DAZAP2 (Supplementary Fig. 6D to G).

Collectively, these findings suggest that DAZAP2 indirectly inhibits SARS-CoV-2 infection, potentially through the regulation of a specific subset of host genes. While the precise mechanisms remain to be fully elucidated, our data highlight DAZAP2 as a multifaceted restriction factor that acts at both the entry and replication stages of the viral life cycle.

### Knockout of *Dazap2* promotes SARS-CoV-2 infection in mouse models

Given the functional conservation of human DAZAP2 and its mouse orthologue in cells-based assays, we extended our investigation to *in vivo* mouse models. To generate the *Dazap2*-knockout mouse (*Dazap2*^-/-^), exons 2 and 3 of the *Dazap2* locus were removed by CRISPR/Cas9 technology, resulting in the deletion of 365 bp coding sequence and disruption of protein function (Fig. 5A). The *Dazap2*^-/-^ mice are viable, fertile and do not exhibit any observable defects.

**Fig. 5.**
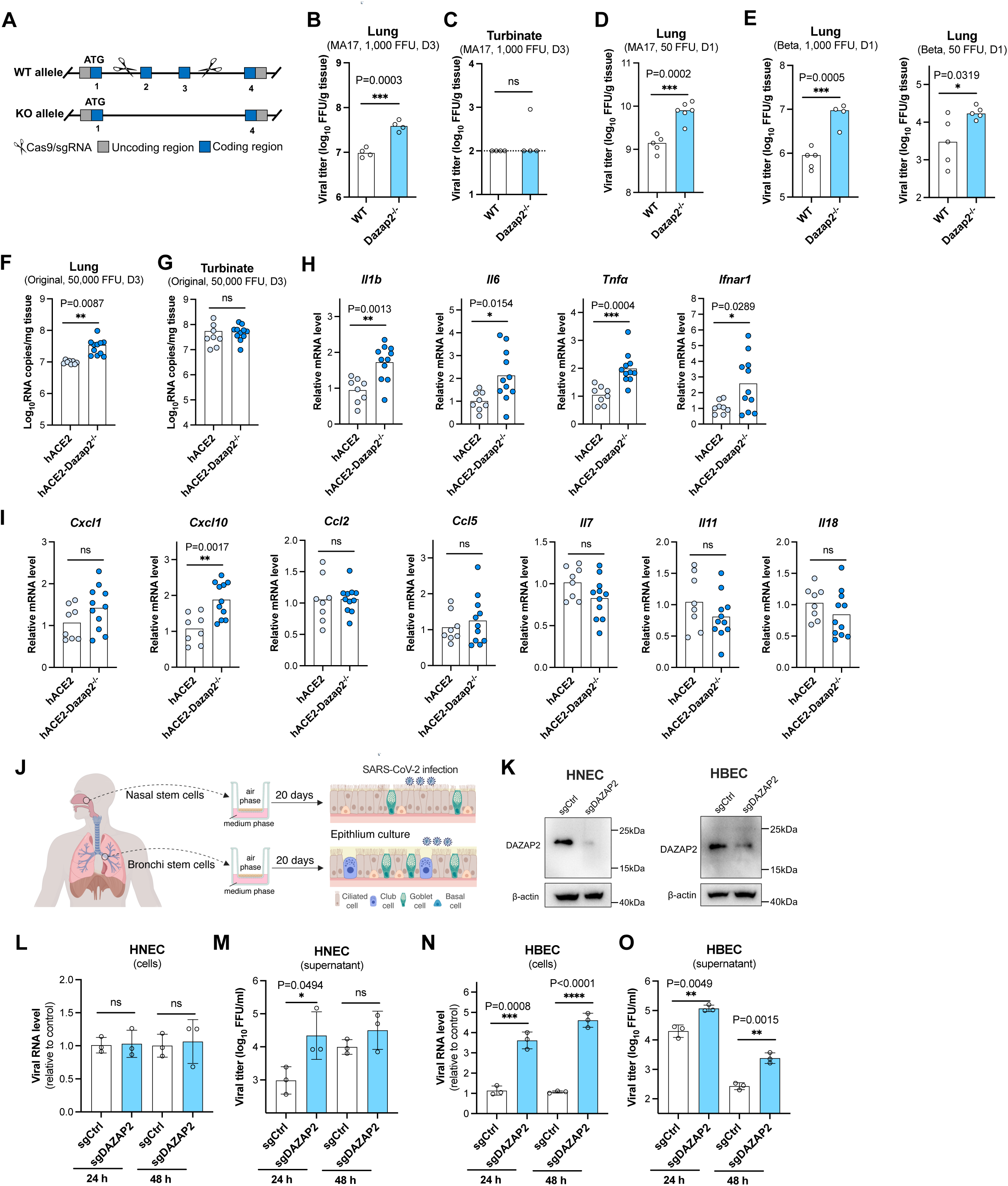
Knockout of *DAZAP2* promotes SARS-CoV-2 infection in mouse models and human primary airway epithelial cells. **A.** Schematic of the generation of *Dazap2-*knockout mice. Exons 2 and 3 were removed using CRISPR/Cas9, resulting in the deletion of 365 bp of coding sequence and disruption of protein function. **B-C.** Female mice at 10-12 weeks old were intranasally inoculated with 1,000 focus-forming unit (FFU) of mouse-adapted Beta variant (B.1.351) of SARS-CoV-2 virus (MA17), and viral loads in the lungs (**B**) or nasal turbinates (**C**) at day 3 post-infection were titrated by focus-forming assay. The dashed line represents the limit of detection. **D.** Mice infected with of 50 FFU of MA17 virus, and viral loads in the lungs at day 1 post-infection were titrated. **E.** Mice infected with 1,000 FFU (left) or 50 FFU (right) of non-adapted Beta variant (B.1.351) of SARS-CoV-2, and viral loads in the lungs at day 1 post-infection were titrated. D-E, one independent experiment with a total of 4-5 mice were used. **F-G.** Human *ACE2* knock-in (h*ACE2*) and h*ACE2* female mice with *Dazap2* deletion (h*ACE2*-*Dazap2*^-/-^) at 10-12 weeks old were intranasally inoculated with 50,000 FFU of original SH01 strain of SARS-CoV-2, and viral loads in the lungs (F) or nasal turbinates (G) were titrated at day 3 post-infection by qRT-PCR. F-G, two independent experiments with a total of 8-11 mice were used. **H-I.** mRNA detection of cytokines in the lungs harvested at day 3 post-infection from F. **J.** Schematic of air-liquid interface cultures of human primary nasal and bronchial epithelial cells for SARS-CoV-2 infection. The figure was created with BioRender.com. **K.** Editing efficiency of DAZAP2 in undifferentiated human nasal epithelial cells (HNEC) and human nasal epithelial cells (HNEC) was validated by western blotting. **L-O.** SARS-CoV-2 infection in differentiated HNEC and HBEC. Viral RNA in cells was determined by qRT-PCR (L and N) and virus production in the supernatant was titrated by focus-forming assay (M and O). Unpaired two-tailed t test; n=3; mean ± s.d. *P < 0.05; **P < 0.01; ***P < 0.001; ****P < 0.0001; ns, not significant.

To assess viral replication, four female age-mapped wild-type (WT) and *Dazap2*^-/-^ mice were intranasally inoculated with the mouse-adapted Beta variant (B.1.351) of SARS-CoV-2. After 3 days, lungs and nasal turbinates were collected for infectious virus titration. While the average titer in the lungs of WT mice reached 7 logs, the average titer in *Dazap2*^-/-^ mice was 7.6 logs (Fig. 5B). Viral titers in the nasal turbinates of both groups were below the limit of detection (Fig. 5C). To further characterize the virus infection in lung tissues, mice were intranasally inoculated with a low dose of 50 FFU of the adapted virus for one day. The average viral titer in *Dazap2*^-/-^ mice increased from 9.15 logs in WT mice to 9.9 logs (Fig. 5D).

The clinical isolate of the Beta variant (B.1.351) of SARS-CoV-2 can naturally infect WT mice, albeit less efficiently than in ACE2 transgenic mice^47–50^. To further evaluate the restriction function of Dazap2, WT and *Dazap2*^-/-^ mice were intranasally challenged with 1,000 or 50 FFU of the non-adapted Beta variant for one day. At higher does (1,000 FFU), viral replication increased from 5.95 logs in WT mice to 6.98 logs in knockout mice (Fig. 5E). A slight increase in viral titer was also observed at the low dose (50 FFU) (Fig. 5E).

To further investigate the restrictive function of DAZAP2, we cross-bred *Dazap2*^-/-^ mice with human ACE2 knock-in mice (hACE2) to enable infection by the original SARS-CoV-2 strain^51^. Female age-matched hACE2 and *Dazap2*^-/-^ mice with ACE2 expression (hACE2-*Dazap2*^-/-^) were intranasally challenged with 50,000 FFU of SH01 strain isolated in early 2020^52^. Viral replication in tissues were quantified by qRT-PCR at day 3 post-infection. Viral titers in the lung of hACE2-*Dazap2*^-/-^ mice were significantly higher than in hACE2 mice (Fig. 5F), while no difference was detected in the nasal turbinates (Fig. 5G). Additionally, the increased viral loads in the lungs of hACE2-*Dazap2*^-/-^ mice were associated with elevated cytokine expression. The levels of *Il1b, Il6, Tnfα, Ifnar1, and Cxcl10* mRNA were significantly higher in hACE2-*Dazap2*^-/-^ mice compared to hACE2 mice (Fig. 5H and I). These results collectively demonstrate that knockout of *Dazap2* enhances SARS-CoV-2 replication in mouse models.

### Knockout of *DAZAP2* promotes SARS-CoV-2 infection in human primary airway epithelial cells

To determine whether DAZAP2 exerts antiviral function in humans, we utilized air-liquid interface (ALI) culture of human primary nasal (HNEC) and bronchial epithelial cells (HBEC). These cultures represent physiologically relevant models for studying SARS-CoV-2 infection (Fig. 5J). Nasal and bronchial stem cells were isolated and edited with control or *DAZAP2*-specific sgRNA, and knockout efficiency was confirmed by western blotting (Fig. 5K).

After differentiation in transwells, the cells were infected with SARS-CoV-2, and viral production was assessed at 24 and 48 hours post-infection. In nasal epithelial cells, we observed a modest 0.74-log increase in virus production in the supernatant of *DAZAP2*-edited nasal epithelial cells at 24 hours (Fig. 5L and M). In contrast, bronchial epithelial cells exhibited significantly higher levels of viral RNA in cells and viral titers in the supernatant at both 24 and 48 hours post-infection in *DAZAP2*-edited cells when compared to control cells (Fig. 5N and O). These findings demonstrate that DAZAP2 acts as a host restriction factor in human airway tissues, significantly inhibiting SARS-CoV-2 infection in physiologically relevant models. The stronger antiviral effect observed in bronchial epithelial cells highlights the importance of DAZAP2 in protecting the lower respiratory tract from viral infection.

## DISCUSSION

The global spread of SARS-CoV-2 has caused unprecedented devastation and mortality since the 1918 influenza pandemic. Identifying the host factors involved in SARS-CoV-2 infection is critical for understanding virus-host interactions, elucidating pathogenesis, and developing potential host-directed therapeutics. While proviral host factors have been extensively studied using high-throughput strategies, such as genome-wide CRISPR screens^16,17,52–58^, antiviral host factors remain less explored. Although several studies have focused on screening of ISGs, a large family of antiviral genes^12–15,18^, a comprehensive genome-scale identification of host restriction factors is still needed.

Unlike genome-wide activation screens^16,17^, or CRISPR dropout screens analyzing depleted cells^29^, we employed a FACS-based genome-wide CRISPR knockout screen to enrich the highly susceptible live cells. Through this work, we identified and validated a set of restriction factors, including *PDCD10*, *DAZAP2*, *CAB39*, and *VTA1*. Among these, the known IFN-related gene *PLSCR1* exhibited the most potent antiviral activity^27,40–42^. We focused on the gene with the second greatest impact, *DAZAP2*, which after being knocked out led to an over 4-fold increase in SARS-CoV-2 infection.

Although Hou et al. previously identified DAZAP2 as an antiviral gene regulating *SERPINE* expression ^29^, its mechanisms of action and *in vivo* relevance remained unclear. In this study, we systematically dissected the stages of viral entry and replication, revealing that: a. DAZAP2 inhibits virion fusion with both endolysosomal and plasma membranes, blocking viral entry; b. DAZAP2 inhibits genomic RNA replication but does not affect the primary translation of replicases from incoming genomes. Notably, the restriction of DAZAP2 against cell entry appears more pronounced than its effect on replication, as demonstrated by pseudovirus and replicon assays, respectively. We also established DAZAP2 as a pan-coronavirus restriction factor, functioning across four genera of coronaviruses, and validated its antiviral roles in mouse models and human primary airway epithelial cells.

DAZAP2 is an evolutionary conserved protein ubiquitously expressed across tissues^59–61^. It contains predicted Src homology 2 (SH2)/SH3 binding sites and a polyproline region, and is reported to localize in both the cytoplasm and nucleus^36–38,59,61^. Its nuclear localization suggests a regulatory role in gene expression. For example, DAZAP2 interacts with the transcription factor SOX6 to regulate the expression of L-type Ca^2+^ channel α_1c_ gene and cardiac differentiation^62^. Similarly, it interacts with the transcription factor TCF-4 to regulate gene expression, possibly by modulating the affinity of its DNA-recognition motif^37^. DAZAP2 is also implicated in diseases, with its downregulation reported in multiple myeloma^59,63,64^. Intriguingly, DAZAP2 occupies p53 response elements to regulate the expression of a subset of genes upon DNA-damaging chemotherapeutic treatment and plays a protective role in cell survival^36^.

Despite inhibiting virion fusion and genomic replication, DAZAP2 primarily localizes to the nucleus, with no detectable presence in the cytosol, plasma membranes, or endolysosomes. This suggests that the restrictive effects of DAZAP2 are likely indirect, mediated through the regulation of specific host genes. While *SERPINE1* was previously identified as a DAZAP2-regulated gene associated with COVID-19 severity^29^, editing *SERPINE1* had no impact on SARS-CoV-2 infection in our study. This implies that other, yet unidentified, genes regulated by DAZAP2 may underlie its antiviral effects. We hypothesize that DAZAP2 interacts with host transcription factors to modulate the expression of specific genes or pathways. Alternatively, *DAZAP2* knockout may disrupt cellular homeostasis, rendering cells more susceptible to viral infection.

Stress granules (SGs) are host organelles that protect cells from harmful stress or virus infection by sequestering host and viral factors, including proteins and RNAs. In addition to its nuclear localization, DAZAP2 is found in the cytoplasm, where it participates in SG formation^38^. DAZAP2 interacts with the RNA-binding protein deleted-in-azoospermia-like (DAZL), which is essential for SG formation and germ cell protection under heat stress^65^. Whether the restrictive effect of DAZAP2 on coronavirus replication involves its role in SGs remains to be determined. Notably, coronaviruses encode proteins such as N protein, NSP1, and NSP15 to disrupt SG formation and evade antiviral responses^66–71^. This may explain why DAZAP2 puncta were not observed in the cytoplasm during infection.

To validate the restriction effect of DAZAP2 on SARS-CoV-2 infection, we generated *Dazap2*-knockout mice. Consistent with previous reports, these mice exhibited no obvious developmental abnormalities^60^. Upon viral challenge, viral loads in the lungs were significantly higher in knockout mice than the WT mice, confirming the protective role of DAZAP2 during infection. Similarly, in air-liquid interface cultures of human primary bronchial epithelial cells, SARS-CoV-2 infection was significantly enhanced in DAZAP2-edited cells when compared to the control cells. Interestingly, no difference in viral titers was observed in the nasal turbinates of mice infected with the Beta variant (B.1.351) or the original strain. Similarly, only minor phenotypic differences were detected in primary nasal epithelial cells. This may reflect the inefficient replication of these viruses in the upper respiratory tract and nasal tissues^72^.

In summary, we conducted a FACS-based genome-wide CRISPR knockout screen to identify SARS-CoV-2 host restriction factors, including the previously known *DAZAP2*. We thoroughly dissected its roles in viral entry and replication and validated its antiviral functions in mouse models and human primary airway epithelial cells. These findings provide new insights into the host defense system against coronavirus infection and highlight potential avenues for developing host-directed therapeutics.

## METHODS

### Cells and viruses

Vero E6 (Cell Bank of the Chinese Academy of Sciences, Shanghai, China), HEK 293T (ATCC #CRL-3216), A549 (ATCC #CCL-185), A549-ACE2^52^, HeLa (ATCC #CCL-2), HeLa-ACE2^52^, Calu-3 (Cell Bank of the Chinese Academy of Sciences, Shanghai, China), MEF expressing the human ACE2 (MEF-ACE2), BHK-21, Huh7, swine testicular (ST), LLC-MK2, HRT-18, and 11*MAVS A549*^73^, all were cultured at 37°C in Dulbecco’s Modified Eagle Medium supplemented with 10% fetal bovine serum (FBS), 10 mM HEPES, 1 mM Sodium pyruvate, 1× non-essential amino acids, and 100 U/ml of Penicillin-Streptomycin. The *IRF3-* and *STAT1-*knockout A549 clonal cell lines were generated by transduction of lentivirus expressing individual sgRNA and selected for 7 days with puromycin and blasticidin, respectively. Clonal cell lines were obtained by limiting dilution and verified by western blotting. All cell lines were tested routinely and free of mycoplasma contamination.

The SARS-CoV-2 (nCoV-SH01-Sfull) stock^52^, swine acute diarrhea syndrome coronavirus (SADS-CoV), porcine epidemic diarrhea virus (PEDV), and infectious bronchitis virus (IBV) were propagated in Vero E6 cells and titrated in the Vero E6 by focus-forming assay^74^. Other virus stock of coronaviruses, porcine deltacoronavirus (PDCoV) (ST cells), HCoV-229E (Huh7 cells), HCoV-OC43 (HRT-18 cells), were prepared and titrated similarly in their respective cell lines. All experiments involving SARS-CoV-2 live virus infection were performed in the biosafety level 3 (BSL-3) facility of Fudan University or Guangzhou Customs Technology Center following the regulations.

### Genome-wide CRISPR knockout screen

The human Brunello CRISPR knockout pooled library encompassing 76,441 different sgRNAs targeting 19,114 genes^30^ was a gift from David Root and John Doench (Addgene #73178), and packaged in 293FT cells after co-transfection with psPAX2 (Addgene #12260) and pMD2.G (Addgene #12259) at a ratio of 2:2:1 using Fugene^®^HD (Promega). At 48 h post transfection, supernatants were harvested, clarified by spinning at 3,000 rpm for 15 min, and aliquoted for storage at -80 °C.

For the CRISPR sgRNA screen, A549-ACE2-Cas9 cells^52^ were transduced with packaged sgRNA lentivirus library at a multiplicity of infection (MOI) of ∼0.3 by spinoculation at 1000g and 32 °C for 30 min in 12-well plates. After selection with puromycin for around 7 days, cells were inoculated with SARS-CoV-2 transcription- and replication-competent virus-like particles that the N gene is replaced by the reporter GFP (trVLP-GFP)^31^. trVLP-GFP was packaged in cells expressing the N gene and only replicated for single round in A549-ACE2 in the absence of N protein. After infection at an MOI of 0.5 for 24 h, cells were harvested and sorted for the GFP positive population. Genomic DNA from both sorted cells and uninfected cells was extracted for sgRNA amplification and next generation sequencing using an Illumina NovaSeq 6000 platform. The sgRNA sequences targeting specific genes were trimmed using the FASTX-Toolkit (http://hannonlab.cshl.edu/fastx_toolkit/) and cutadapt 1.8.1, and further analyzed for sgRNA abundance and gene ranking by a published computational tool (MAGeCK) (see Supplementary table 1).

### Gene validation

Top 20 genes from the MAGeCK analysis were selected for validation. Two independent sgRNAs per gene were chosen from the Brunello CRISPR knockout library and cloned into the plasmid lentiCRISPR v2 (Addgene #52961) and packaged with plasmids psPAX2 and pMD2.G. A549-ACE2 cells were transduced with lentiviruses expressing individual sgRNA and selected with puromycin for 7 days. The gene-edited mixed population of cells was used for validation using the SARS-CoV-2 transcription- and replication-competent virus-like particles that the N gene is replaced by the NanoLuc luciferase (trVLP-Nluc) at an MOI of 0.5 for 24 h. trVLP-Nluc was constructed in this study to measure virus replication for convenience. The luciferase activity was determined using Nano-Glo^®^ Luciferase Assay kit (Promega #N1110) and the luminescence was recorded by using a FlexStation 3 (Molecular Devices). The sgRNA sequences are listed in Supplementary table 2.

For authentic SARS-CoV-2 virus infection, gene-edited A549-ACE2, HeLa-ACE2, Calu-3, and MEF-ACE2 cells were inoculated for 24 h. Cells were fixed with 4% paraformaldehyde (PFA) diluted in PBS for 30 min at room temperature, and permeabilized with 0.2% Triton x-100 in PBS for 1 h at room temperature. Cells then were subjected for immunofluorescence staining and high-content imaging. The gene-edited A549-ACE2, HeLa-ACE2, or MEF-ACE2 cells were also infected with members of family *Coronaviridae*, and then subjected for immunofluorescence staining, high-content imaging or flow cytometry analysis.

### Pseudotyped virus experiment

SARS-CoV-2 pseudoviruses were packaged as previously described^52^. Shortly, pcDNA3.1 vector expressing the spike gene of SARS-CoV-2 lacking the C-terminal 21 amino acids, the full spike of SARS-CoV-1, or VSV-G (pMD2.G (Addgene #12259)), was co-transfected in HEK 293T cells with the murine leukemia retrovirus (MLV) expressing the NanoLuc luciferase gene and plasmid expressing the MLV Gag-Pol using Fugene^®^HD transfection reagent (Promega). The virus entry was assessed by transduction of pseudoviruses in gene-edited cells in 96-well plates. After 48 h, the luciferase activity was determined using Nano-Glo^®^ Luciferase Assay kit (Promega #N1110) and the luminescence was recorded by using a FlexStation 3 (Molecular Devices).

### Generation of SARS-CoV-2 replicon system and trVLP-Nluc particles

To construct the replicon system, the full-length viral RNA of SARS-CoV-2 (nCoV-SH01, GenBank accession no. MT121215) was reverse-transcribed, PCR-amplified as 6 fragments, and cloned into pSMART vector individually. The region encompassing the spike gene to ORF8 was replaced by NanoLuc luciferase-P2A-puromycin cassette. The T7 promoter was inserted upstream of the viral genome to initiate the transcription. The sequences of HDVr ribosome and transcription terminater were added downstream of poly-A tail of viral genome. Six pSMART plasmids and pBeloBAC11 vector were digested with type IIS restriction enzymes and assembled in vitro as one plasmid, and then amplified in bacterium. Replicon RNA was transcribed from the single plasmid using mMESSAGE mMACHINE T7 Transcription Kit (Invitrogen #AM1344) according to the manufacturer’s instructions. RNA was then transfected into target cells to assess the virus replication efficiency by measuring the luciferase activity using Nano-Glo^®^ Luciferase Assay kit (Promega #N1110).

To detect the virus infection conveniently and safely, SARS-CoV-2 transcription- and replication-competent virus-like particles that the N gene is replaced by the NanoLuc luciferase (trVLP-Nluc) were generated as described previously^31^. The replicon plasmid constructed above was modified by maintaining the spike to ORF8 genes but replacing the N gene with NanoLuc luciferase. The single round trVLP-Nluc particles was packaged in Vero E6 cells expressing the N gene. The virus infection was determined by measuring the luciferase activity using Nano-Glo^®^ Luciferase Assay kit (Promega #N1110).

To modify the replicon system to assess the primary translation of genomic RNA, the NanoLuc luciferase-P2A-puromycin cassette in the replicon constructed above was removed, and a coding sequence of P2A-NanoLuc luciferase cassette was inserted at the NSP1/NSP2 junction. To make a replication-deficient replicon, the viral RdRp mutant was created by mutating NSP12 catalytic residues at positions 760 and 761 from aspartic acid (D) to asparagine (N)^46^. As described above, modified replicon RNA was *in vitro* transcribed and then electroporated into target cells to assess the primary translation of genomic RNA by measuring the luciferase activity.

### Plasmid constructs

The human DAZAP2 (Sino Biological #HG15906-G) or mouse Dazap2 (Sino Biological #MG52462-G) gene was PCR-amplified and cloned into the pLV-EF1α-IRES-puro (Addgene #85132). The PH-Halo-LgBiT fragment was synthesized (GENEWIZ), amplified, and cloned into pLV-EF1α-IRES-Hygro. The LgBiT fragment was cloned into pCAGGS vector by using the synthesized PH-Halo-LgBiT as template. The CypA-HiBiT fragment was synthesized (GENEWIZ), amplified, and cloned into pCAGGS. Lentiviruses were packaged by co-transfection with psPAX2 (Addgene #12260) and pMD2.G (Addgene #12259) and transduced into wildtype, DAZAP2-deficient A549-ACE2, or MEF-ACE2 cells.

### Protein lysate preparation and western blotting

Cells in plates washed twice with ice-cold PBS and lysed in RIPA buffer (Cell Signaling #9806S) with a cocktail of protease inhibitors (Sigma-Aldrich #S8830). Samples were prepared in reducing buffer (50 mM Tris, pH 6.8, 10% glycerol, 2% SDS, 0.02% [wt/vol] bromophenol blue, 100 mM DTT). After heating (95°C, 10 min), samples were electrophoresed in 10% SDS polyacrylamide gels, and proteins were transferred to PVDF membranes. Membranes were blocked with 5% non-fat dry powdered milk in TBST (100mM NaCl, 10mM Tris, pH7.6, 0.1% Tween 20) for 1 h at room temperature, and probed with the primary antibodies at 4 °C overnight. After washing with TBST, blots were incubated with horseradish peroxidase (HRP)-conjugated secondary antibodies for 1 h at room temperature, washed again with TBST, and developed using SuperSignal West Pico or Femto chemiluminescent substrate according to the manufacturer’s instructions (Thermo fisher).

The antibodies used are as follows: mouse anti-DAZAP2 (Santa Cruz #sc-515182, 1:1000), mouse anti-FLAG (Sigma #F1804, 1:2000), mouse anti-HA (Abmart #M20003M, 1:2000), mouse anti-FURIN (Proteintech #67481-1-Ig, 1:2000), rabbit anti-TMPRSS2 (Abcam #ab109131, 1:1000), mouse anti-CTSL (Thermo fisher #BMS1032, 1:1000), rabbit anti-beta actin (Proteintech #20536-1-AP, 1:5000), mouse anti-GAPDH (Proteintech #60004-1-Ig, 1:2000), home-made mouse serum against N protein from SARS-CoV-2, HCoV-229E, PEDV, SADS-CoV or PDCoV (1:1000), rabbit anti-HCoV-OC43 N (Sino Biological #40643-T62, 1:1000), home-made mouse anti-MHV N mAb (1:1000), rabbit anti-HCoV-NL63 N (Sino Biological #40641-T62, 1:1000), home-made mouse anti-IBV N mAb (1:1000). The HRP-conjugated secondary antibodies include: Goat anti-mouse (Sigma #A4416, 1:5000), goat anti-rabbit (Thermo fisher #31460, 1:5000).

### Immunofluorescence staining and analysis

For high-content imaging analysis, virus-infected cells in plates were fixed with 4% paraformaldehyde in PBS for 30 min, permeablized with 0.2% Triton X-100 for 1 h. Cells were then incubated with house-made mouse serum (1:1000) against nucleocapsid protein from different coronaviruses for 2 h at room temperature. After three washes, cells were incubated with the secondary goat anti-mouse IgG (H + L) conjugated with Alexa Fluor 555 (Thermo fisher #A-21424, 2 μg/ml) for 1 h at room temperature, followed by staining with 4’,6-diamidino-2-phenylindole (DAPI). Images were collected using an Operetta High Content Imaging System (PerkinElmer), and processed using the PerkinElmer Harmony high-content analysis software v4.9 and ImageJ v2.0.0 (http://rsb.info.nih.gov/ij/).

For flow cytometry analysis, virus-infected cells were harvested with trypsin, and fixed with 2% paraformaldehyde in PBS for 10 min. Cells were permeablized with 0.1% saponin in PBS for 10 min, and stained with house-made mouse serum (1:1000) against nucleocapsid protein from different coronaviruses for 30 min at room temperature. After washing, cells were incubated with the secondary goat anti-mouse IgG (H + L) conjugated with Alexa Fluor 647 (Thermo fisher #A21235, 2 μg/ml) for 30 min at room temperature. After two additional washes, cells were subjected to flow cytometry analysis (Thermo, Attune™ NxT) and data processing (FlowJo v10.0.7).

For surface staining of entry-related host factors and analyzed by flow cytometry, cells were collected with TrypLE (Thermo fisher #12605010) and incubated with the rabbit anti-ACE2 (Sino Biological #10108-RP01, 1:500), mouse anti-AXL (R&D #MAB154-SP, 1 μg/ml), mouse anti-DC-SIGN (Biolegend #330102, 1 μg/ml), mouse anti-TIM-1 (Biolegend #354002, 1 μg/ml), mouse anti-SIGLEC1 (Abcam #ab199401, 1 μg/ml), or mouse anti-heparan sulfate (10E4) (USBiological #H1890, 1 μg/ml) primary antibody at 4 °C for 30 min. After washing, cells were stained with goat anti-rabbit IgG (H + L) conjugated with Alexa Fluor 647 (Thermo fisher #A21245, 2 μg/ml) or goat anti-mouse IgG (H + L) conjugated with Alexa Fluor 647 (Thermo fisher #A21235, 2 μg/ml) for 30 min at 4 °C and subjected to flow cytometry analysis.

For confocal microscopy analysis, cells seeded on coverslips were fixed with 4% paraformaldehyde in PBS for 30 min, permeablized with 0.1% saponin in PBS for 10 min. Cells were then incubated with primary antibody overnight at 4 °C. After three washes, cells were incubated with the secondary antibody for 2 h at room temperature, followed by staining with 4’,6-diamidino-2-phenylindole (DAPI). Images were collected using Leica Confocal Microscope (TCS SP8), processed using the Leica Application Suite X (LAS X, v3.7.0.20979) and ImageJ v2.0.0 (http://rsb.info.nih.gov/ij/). The primary antibodies used as follows: house-made mouse anti-SARS-CoV-2 nucleocapsid protein serum (1:1000), rabbit anti-SARS-CoV-2 spike (Sino Biological #40591-T62, 1:1000), rabbit anti-LAMP1 (Abcam #ab24170, 1:1000), mouse anti-DAZAP2 (Santa Cruz #sc-515182, 1:1000), rabbit anti-HA (Abcam #ab9110, 1:1000), mouse anti-dsRNA antibody (J2) (Scicons #10010200). The secondary antibodies used as follows: goat anti-mouse IgG (H + L) conjugated with Alexa Fluor 555 (Thermo fisher #A-21424, 2 μg/ml), goat anti-rabbit IgG (H + L) conjugated with Alexa Fluor 488 (Thermo fisher #A-11034, 2 μg/ml), followed by staining with 4’,6-diamidino-2-phenylindole (DAPI).

### Virus binding and internalization assay

For binding assay, A549-ACE2 cells were pre-chilled on ice for 10 min followed by incubation with ice-cold virus (MOI 5) on ice for 45 min. After washing with ice-cold PBS three times, cells were lysed in TRIzol reagent (Thermo fisher #15596018) for RNA extraction and qRT-PCR.

For internalization assay, after virus binding as described above, cells were washed with ice-cold PBS three times, followed by incubation at 37 °C for 45 min. Uninternalized virions on the cell surface were removed by treating cells with 400 μg/ml protease K on ice for 45 min. After washing with ice-cold PBS three times, cells were lysed in TRIzol reagent for RNA extraction and qRT-PCR. The relative amount of bound or internalized virions was normalized to internal control GAPDH.

### Virion trafficking assay

The experiments were conducted as described previously^42^. Control or DAZAP2-deficient A549-ACE2 cells seeded on coverslips were pretreated with 25 μM of E-64d, a cathepsin B and L proteinase inhibitor. One hour later, cells were inoculated with SARS-CoV-2 trVLP-Nluc. The inhibitor E-64d was maintained in the medium during the infection. At 4 h post infection, cells were washed twice with PBS, fixed with 4% PFA for 10 min and then permeabilized with 0.1% saponin for 10 min. Cells were blocked with 5% BSA in PBS for 1 h and incubated with primary antibodies (rabbit anti-SARS-CoV-2 spike protein, mouse anti-SARS-CoV-2 nucleocapsid protein or rabbit anti-LAMP1) at 4 °C overnight. After three washes, cells were incubated with the secondary goat anti-mouse or rabbit antibody conjugated with Alexa Fluor 555 or 488 for 2 h at room temperature, followed by staining with DAPI. Images were acquired using Leica Confocal Microscope (TCS SP8), and processed using the Leica Application Suite X (LAS X, v3.7.0.20979). The number of spike and nucleocapsid double-positive particles per field was quantified.

### Quantification of endosomal acidification

Control or *DAZAP2*-deficient A549-ACE2 cells seeded in 96-well plate were pre-treated with or without chloroquine (CQ) (20 μM). One hour later, cells were incubated in phenol red free DMEM containing 2 μM LysoSensor Green dye (Thermo #L7535) in the presence or absence of CQ (20 μM) for 30 min at 37 °C. Images were obtained using an AMG microscope (EVOS M7000) and the fluorescence intensity of LysoSensor were analyzed using ImageJ v2.0.0 (http://rsb.info.nih.gov/ij/).

### Virus-cell fusion assay

The experiments were conducted via an improved system as described previously^42^. The Gag interacting protein cyclophilin A (CypA) was fused with HiBit and encapsulated into MLV retrovirus particles bearing the SARS-CoV-2 spike protein. Pseudoviruses containing CypA-HiBiT were packaged in HEK 293T cells by co-transfecting the retrovector pMIG for which the target gene was replaced by mGreenLantern^52^, plasmid expressing the MLV Gag-Pol, pCAGGS expressing SARS-CoV-2 spike protein with the deletion of C-terminal 21 amino acids, and pCAGGS expressing CypA-HiBiT using Fugene^®^HD transfection reagent (Promega). At 48 h post transfection, the supernatant was harvested, clarified by spinning at 3500 r.p.m. for 15 min, aliquoted, and stored at -80 °C. Control and DAZAP2-deficient A549-ACE2 cells (target cells) were transduced with pLV-PH-Halo-LgBiT-hygro lentivirus to stably express the LgBiT fragment. Target cells were seeded in black/clear bottom 96-well plate for 24 h, followed by spinfection with 50 μl of pseudoviruses per well at 1,000*g*, 4 °C for 30 min. The luciferase activity was determined using Nano-Glo Luciferase Assay kit at 8 h post infection, and the luminescence was recorded by using a FlexStation 3 (Molecular Devices).

### Cell-cell fusion assay

For visualization and quantification of the syncytia formed after cell–cell fusion, HEK293T cells (donor cells) were transfected with pCAGGS vector expressing the spike protein of SARS-CoV-2 that lacks the C-terminal 21 amino acids. At 24 h post transfection, control and DAZAP2-deficient A549-ACE2 cells (acceptor cells) and donor cells were trypsinized and seeded in 24-well plate at a ratio 1:1. After 6 hours of co-culture, cells were washed with PBS and fixed by 4% PFA. Images were obtained using an AMG microscope (EVOS M7000). Cell–cell fusion was quantified by a Wright-Giemsa staining according to the manufacturer’s instructions (Sangon #E607315). Images were obtained using an AMG microscope (EVOS M7000) and the number of syncytia or syncytial nuclei was analyzed using ImageJ v2.0.0 (http://rsb.info.nih.gov/ij/).

To quantify the cell-cell fusion based on luciferase activity as previously reported^42^, control and DAZAP2-edited A549 cells (acceptor cells) were transfected with pCAGGS-LgBiT encoding the LgBiT fragment of the split-NanoLuc luciferase. HEK 293T cells (donor cells) were transfected with pCAGGS-HiBiT encoding the HiBiT fragment of split-NanoLuc luciferase, together with pCAGGS vector expressing the spike protein of SARS-CoV-2 that lacks the C-terminal 21 amino acids. At 24 h post transfection, acceptor and donor cells were trypsinized and seeded together in black/clear bottom 96-well plate at a ratio 1:1. After 24 hours of co-culture, the luciferase activity was determined using Nano-Glo Luciferase Assay kit and the luminescence was recorded by using a FlexStation 3 (Molecular Devices).

### Generation of Dazap2 knockout mice

The mouse *Dazap2* gene is located in chromosome 15 and the sequence of *Dazap2* locus (NC_000081.6) was obtained from NCBI. It has four transcripts with one encoding the protein. The CRISPR/Cas9 technology was employed to remove the exon2 and exon 3, resulting in the deletion of 365 bp coding sequences and disruption of protein function. sgRNA was transcribed in vitro. Cas9 and sgRNA were microinjected into the fertilized eggs of C57BL/6JGpt mice. Fertilized eggs were transplanted to obtain positive F0 mice which were confirmed by PCR and sequencing. A stable F1 generation mouse model was obtained by mating positive F0 generation mice with C57BL/6J mice. The *Dazap2*-knockout (Dazap2^-/-^) mice are viable, fertile and do not exhibit any observable defects. The generation of knockout mice was accomplished with the help of Cyagen Biosciences (Suzhou, China).

### Mouse experiments

The mouse experiment protocol has been approved by the Animal Ethics Committee of School of Basic Medical Sciences at Fudan University (No.20220228-122) with formal written consent. Wildtype and *Dazap2*-knockout female mice in the same background of C57BL/6J at 10-12 weeks old were used in the study. The clinical isolate of Beta variant (B.1.351) of SARS-CoV-2 was passaged for 17 times (MA17) in 6-8 weeks old BALB/c mice to allow adaptation. Both BALB/c and C57BL/6 mice are vulnerable to the infection of mouse-adapted MA17 virus. The experiment protocol has been approved by the Institutional Review Boards of the First Affiliated Hospital of Guangzhou Medical University, and conducted in the BSL-3 facility following the regulations. The mice were inoculated intranasally with 1,000 or 50 focus-forming unit (FFU) of mouse-adapted or wildtype SARS-CoV-2 Beta variant (B.1.351) in a volume of 50 μl. The *Dazap2*^-/-^ mice also cross-breeded with human ACE2 knock-in mice (hACE2) that the mouse ACE2 coding sequences with the exception of signal peptide region are replaced with human ACE2 (Cyagen Biosciences #C001191). The 10-12 weeks old hACE2 or *Dazap2*^-/-^ female mice with human ACE2 expression (hACE2-*Dazap2*^-/-^) were challenged with 50,000 FFU of SH01 strain isolated in early 2020^52^. Mice were euthanized at day 1 or 3 post infection, and the turbinate and lungs were harvested, homogenized in PBS for live virus titration by focus-forming assay. The cytokine production in lungs at day 3 was detected by qRT-PCR after RNA extraction as described below.

### Preparation of HNEC and HBEC *in vitro* models

Human primary nasal stem cells and bronchial stem cells were isolated from nasal swab and bronchoscopic brushing biopsies, donated by lesion-free individuals. Following 20 min digestion with collagenase IV (Gibco #17104019, 1 mg/ml) in Ham’s F12 (Gibco #11765054), dissociated cells were washed thoroughly in cold wash buffer (Ham’s F12, 5% FBS, 100 μg/ml Penicillin-Streptomycin, 100 μg/ml Gentamycin, and 0.25 μg/ml Fungizone) and selectively expanded in a 3T3-J2 culture system, under the protocol as previously described^75^. To knock out *DAZAP2* gene, *DAZAP2*-specific sgRNA or non-targeting sgRNA was cloned into plasmid lentiCRISPR v2 for which the puromycin resistance gene was replaced by turboGFP reporter. Cells were transduced with lentivirus packaged with helper plasmids psPAX2 and pMD2.G, and sorted for GFP-positive cells. The knockout efficiency was confirmed by western blotting. Cells were then subjected to *in vitro* differentiation via air-liquid interface (ALI) modeling. Upon 20-day differentiation in PneumaCult-ALI medium (StemCell Technologies #05001), mature epithelial structures of HNEC and HBEC were histochemically confirmed and proceeded to subsequent infection studies. The protocol of human biopsy acquisition and culture was approved by the Ethics Committees of Shanghai Jiao Tong University Affiliated Sixth People’s Hospital (Approval number: 2017– 090).

For SARS-CoV-2 infection, differentiated cells on insert of 6.5 mm transwells were incubated with 100 μl of culture medium containing 40,000 FFU of virus for 1 h, and washed for 3 times. At 24 or 48 h post infection, 200 μl of culture medium was added onto cells. After 30 min of incubation, the medium was harvested for FFA, and the cells were lysed for qRT-PCR to determine the virus replication.

### qRT-PCR

RNA from tissues or cells was extracted with the TRIzol reagent (Thermo fisher #15596018). Host mRNAs were determined using the One Step PrimeScript™ RT-PCR Kit (TaKaRa #RR064B) on CFX Connect Real-Time System (Bio-Rad) instrument. Host mRNAs was also reverse transcribed into cDNA with PrimeScript™ RT Reagent Kit (TaKaRa #RR047A) using RT primer mix of Oligo dT primer and random 6 mers. qPCR was performed using TB Green® Premix Ex Taq™ II (TaKaRa #RR820A) on the CFX Connect Real-Time System (Bio-Rad) instrument. Relative gene expression was calculated as relative to GAPDH. Primers used for qRT-PCR are listed in Supplementary table 3.

### Statistical analysis

Statistical significance was assigned when *P* values were < 0.05 using Prism Version 9 (GraphPad). Data analysis was determined by an ANOVA, unpaired t-test, or Mann-Whitney depending on data distribution and the number of comparison groups.

## DATA AVAILABILITY

The authors declare that all relevant data supporting the findings of this study are available within the paper and its Supplementary information. The Supplemental Data provide information for the CRISPR screen, qRT-PCR, and RNAseq analysis. Source data are provided with this paper. Any other data of this study are available upon request.

## ACKNOWLEDGEMENTS

Grants from the National Natural Science Foundation of China (32270163 to R.Z., 32041005 to R.Z., 82341084 to Q.D., 81971500 to J.Z., 92169110 to C.L.), National Key Research and Development Program of China (2024YFC2607300 and 2020YFA0707701 to R.Z.), Program of Shanghai Academic/Technology Research Leader (22XD1420600 to R.Z.), Shanghai Municipal Science and Technology Major Project (ZD2021CY001), Non-profit Central Research Institute Fund of Chinese Academy of Medical Sciences (2023- PT310-02), Shenzhen Medical Research Fund (SMRF No. B2302029), Foundation of Key Laboratory of Structural Biology of Zhejiang Province of Westlake University (to R.Z.), and Natural Science Foundation of Shanghai (19ZR1470400 to X.H.) supported this work. We wish to acknowledge Xiaoqing Sun, Yao Wang, and Shen Cai at Key Laboratory of Medical Molecular Virology (MOE/NHC/CAMS), Shanghai Frontiers Science Center of Pathogenic Microorganisms and Infection, School of Basic Medical Sciences of Fudan University for the help with next generation sequencing, flow cytometry, and imaging analysis, respectively. We thank colleagues at the Biosafety Level 3 Laboratory of Fudan University for help with the technical assistance.

## AUTHOR CONTRIBUTIONS

F.F., R.L., J.C., Y.Z., Y.M., Z.W., Y.W., Z.G., L.Y., Y.Y., R.Z. performed the experiments. F.F., Y.Z., R.Z. designed the experiments. Y.Liu, Y.Sun, Y.Liao, X.H., Q.Z., Y.Huang, L.Q., J.W., Jingxian Zhao, C.L. provided technical or material support. Q.D., Y.X., Z.Y., Y.Hong, P.Z., J.S., Jincun Zhao, R.Z. provided administrative, supervision support. F.F., Z.G., R.Z. performed data analysis. R.Z. wrote the initial draft of the manuscript, with the other authors contributing to editing into the final form.

## COMPETING INTERESTS

The authors declare no competing interests.

## SUPPLEMENTARY FIGURES LEGENDS

**Supplementary Figure 1.**
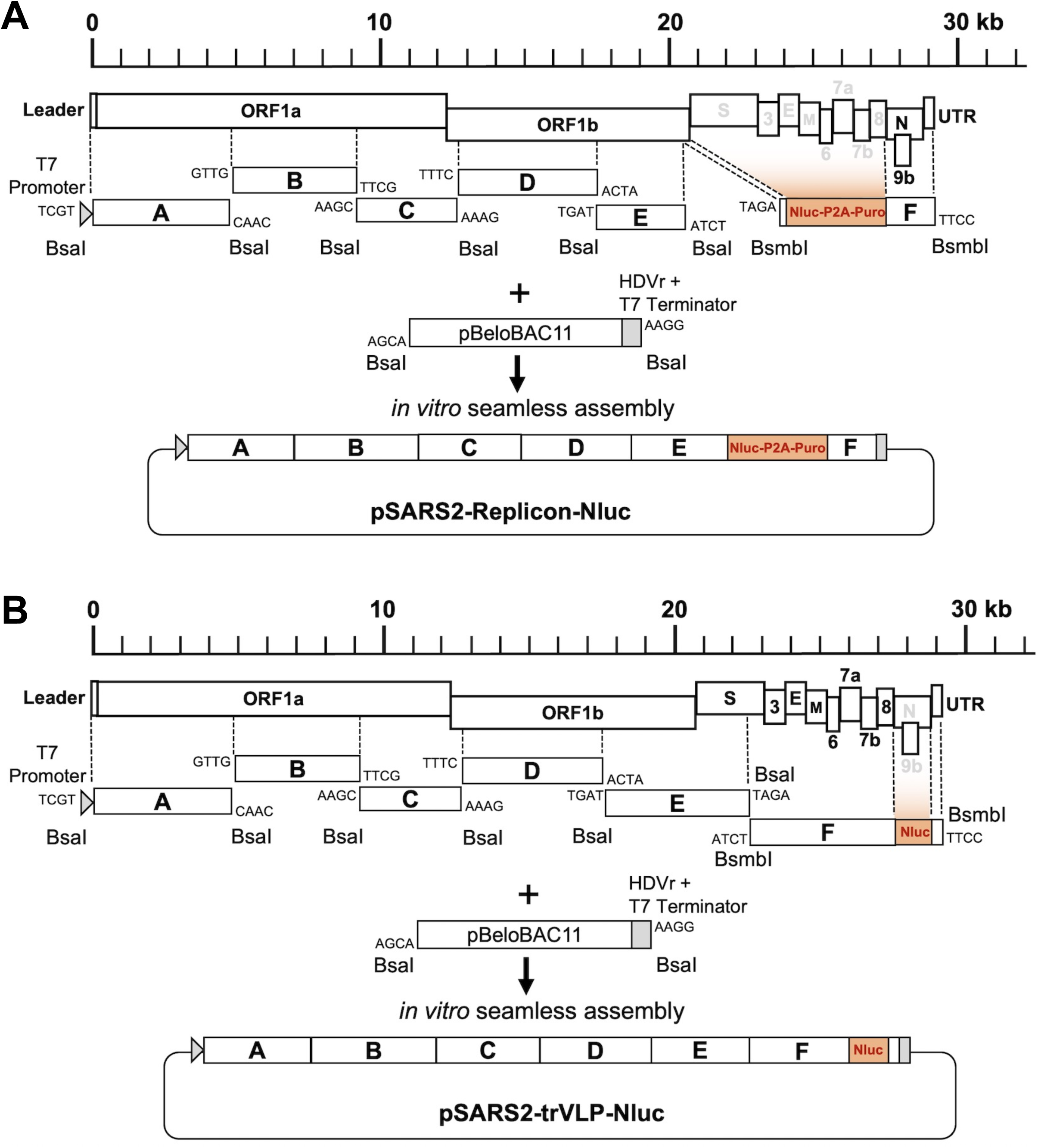
Generation of SARS-CoV-2 replicon system and trVLP-Nluc particles. **A.** Schematic of the construction of SARSC-CoV-2 replicon system. The genome of SARS-CoV-2 was divided into 6 fragments, amplified, and cloned into pSMART vectors. The genes from spike to ORF8 were replaced by a NanoLuc luciferase-P2A-puromycin cassette. Viral fragments were cleaved from the vectors and assembled with the linearized pBeleBAC11 vector *in vitro* using type IIS restriction enzymes. The transcription of replicon RNA was initiated by a T7 promoter and terminated by a T7 terminator. The HDVr sequences were added after the poly-A tail to obtain the correct viral RNA. **B.** Schematic of the generation of trVLP-Nluc particles. Based on the replicon system constructed above, the spike to ORF8 genes were maintained but the N gene was replaced by NanoLuc luciferase. The trVLP-Nluc particles were packaged in Vero E6 cells expressing the N gene, and could only replicate for a single round in cells without the expression of N protein.

**Supplementary Figure 2.**
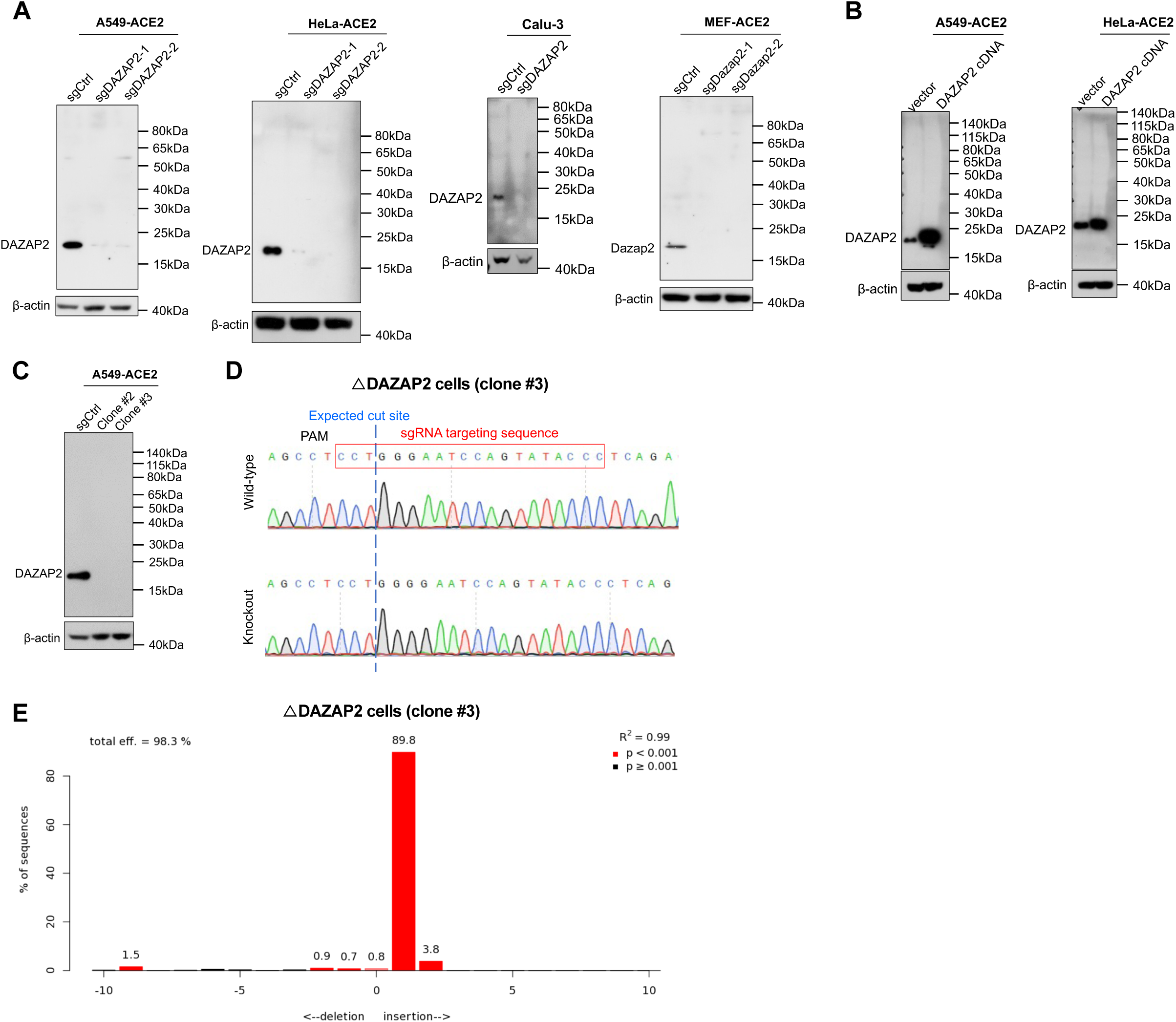
Verification of DAZAP2 expression in cDNA-overexpressing, sgRNA-edited bulk or clonal cells. **A.** Knockout efficiency of *DAZAP2* in A549-ACE2, HeLa-ACE2, Calu-3, and MEF-ACE2 cells. Two sgRNAs or one representative sgRNA were used, and gene-edited bulk cells were subjected to western blotting. **B.** Overexpression of DAZAP2 in A549-ACE2 or HeLa-ACE2 cells. Cells were transduced with lentivirus bearing the human DAZAP2 cDNA, and subjected to western blotting. **C.** Western blotting to verify the *DAZAP2*-knockout clones #2 and #3 of A549-ACE2. Clone #3 (ΔDAZAP2) was used in this study. **D.** The sequence traces of the *DAZAP2* gene locus of WT and clonal cells. The sgRNA target site is indicated. **E.** The ICE analysis of the ΔDAZAP2 clonal cell line.

**Supplementary Figure 3.**
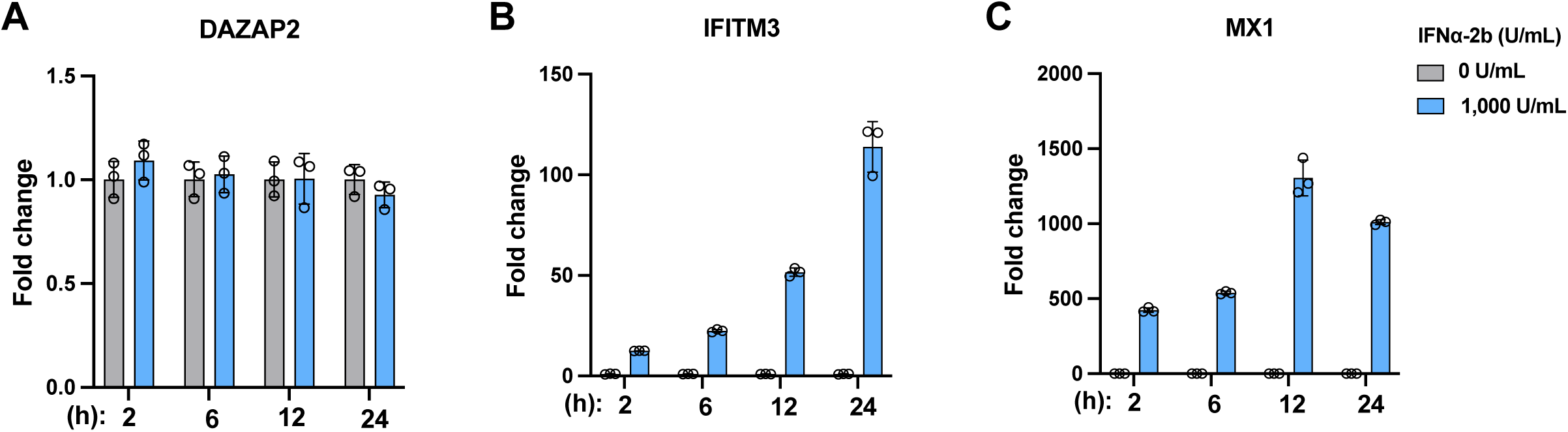
DAZAP2 is not an interferon-stimulated gene (ISG). **A-C.** A549-ACE2 cells were treated with 0 or 1,000 U/ml of IFNα-2b for various time points, and cellular RNA was extracted for detection of *DAZAP2* (**A**), *IFITM3* (**C**), or *MX1* (**C**) by qRT-PCR. The experiment was performed in triplicate and data were normalized to the treatment with 0 U/ml of IFNα-2b at each time point.

**Supplementary Figure 4.**
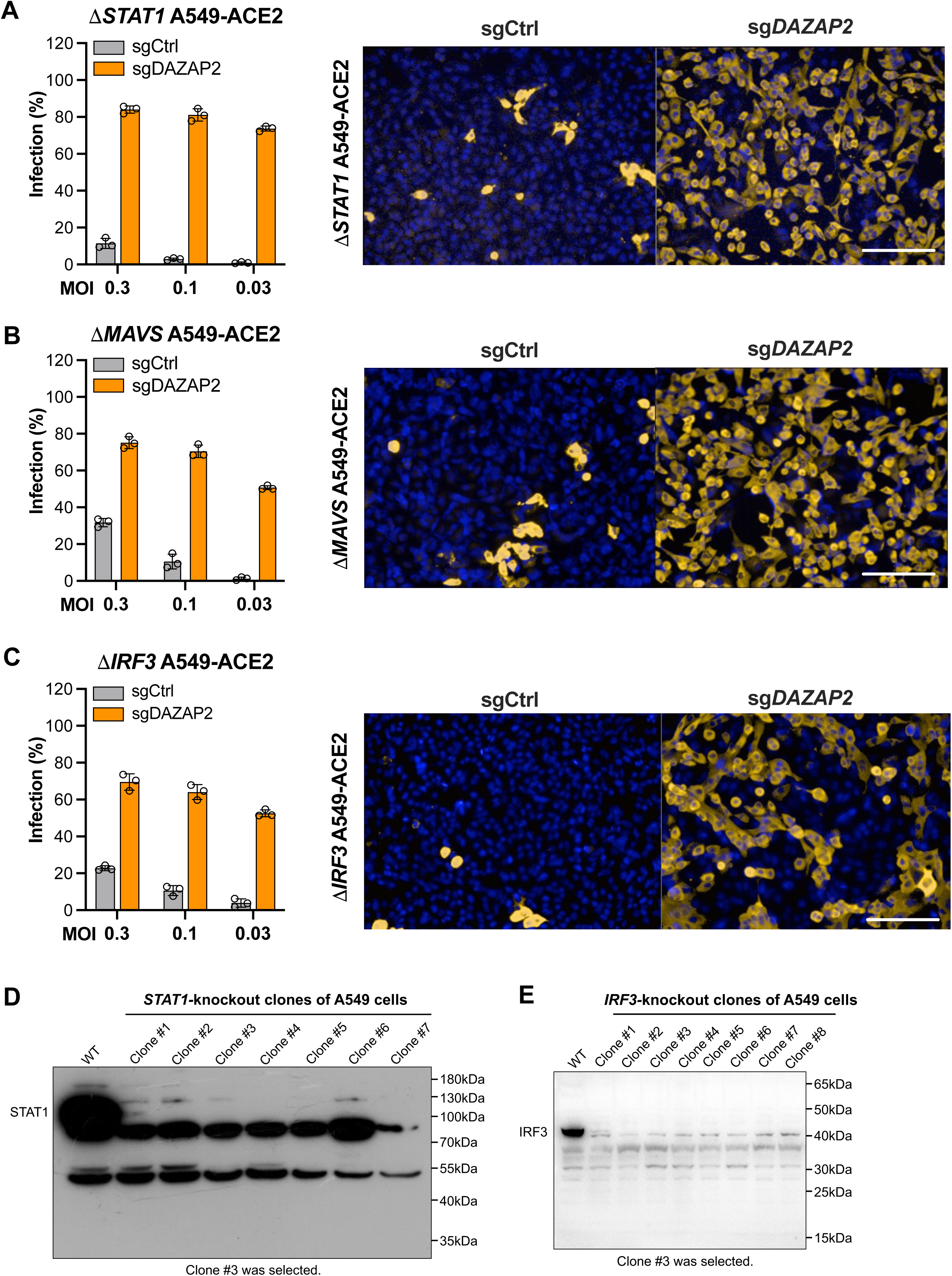
The antiviral effect of DAZAP2 in innate immune gene-knockout cells. **A-C.** The antiviral effect of DAZAP2 on SARS-CoV-2 infection in *STAT1*-, *MAVS*-, or *IRF3*-knockout A549-ACE2 cells. Cells were infected with different MOIs and the percentage of N positive cells were analyzed (left panels of A to C). The representative fluorescence images from high-content analysis in cells were shown on right panels of A to C. **D-E.** Verification of *STAT1-* or *IRF3-*knockout efficiency in clonal cell lines by Western blotting. The clone #3 for *STAT1-* or *IRF3-*knockout cells was selected for use.

**Supplementary Figure 5.**
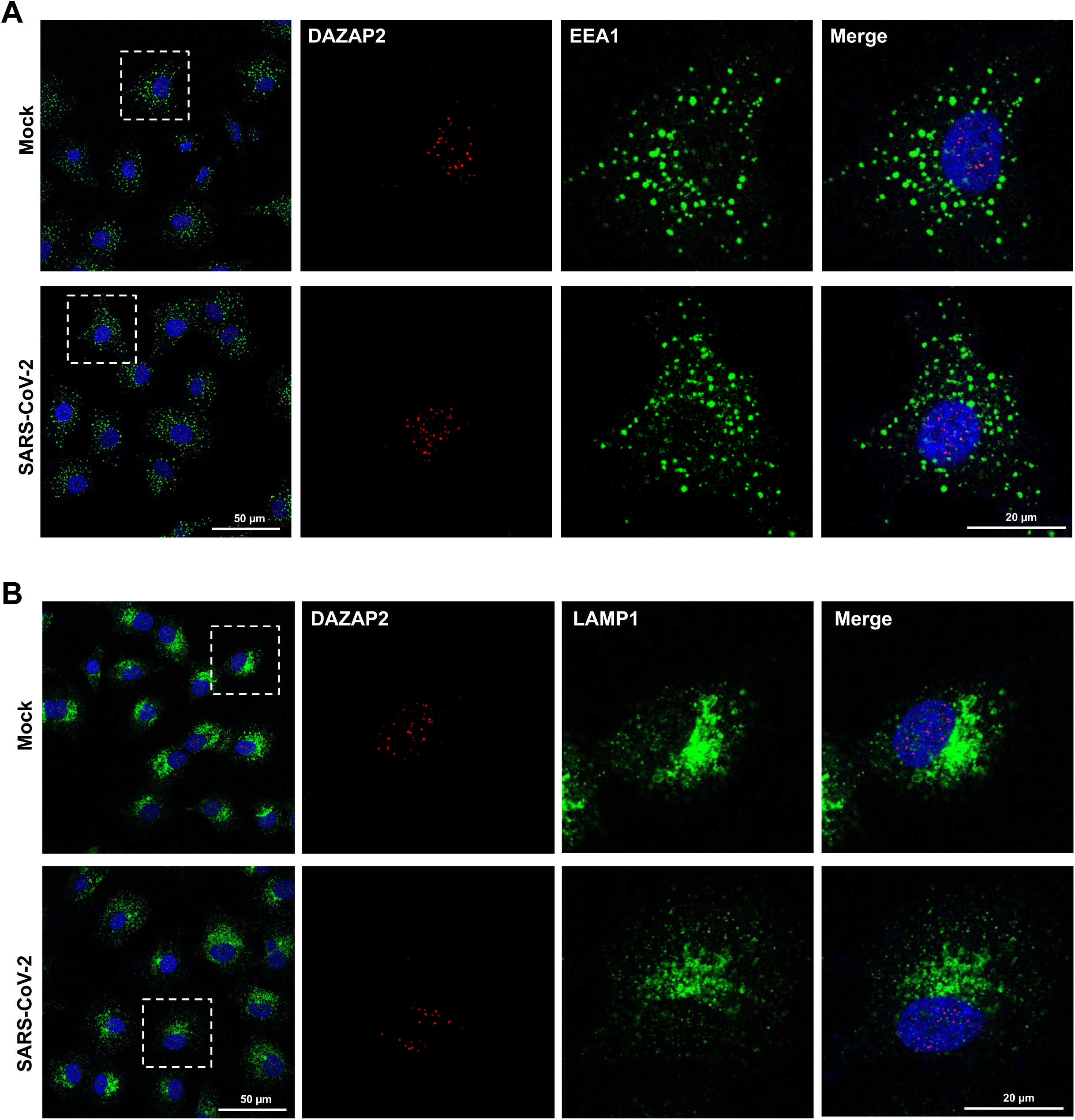
Localization study of DAZAP2 by confocal microscopy. **A-B.** The A549-ACE2 cells were infected with SARS-CoV-2, then fixed and stained with anti-DAZAP2 (A and B), anti-EEA1 (A), or anti-LAMP1 (B) antibody. The representative confocal images were shown. Scale bar, 50 or 20 μm.

**Supplementary Figure 6.**
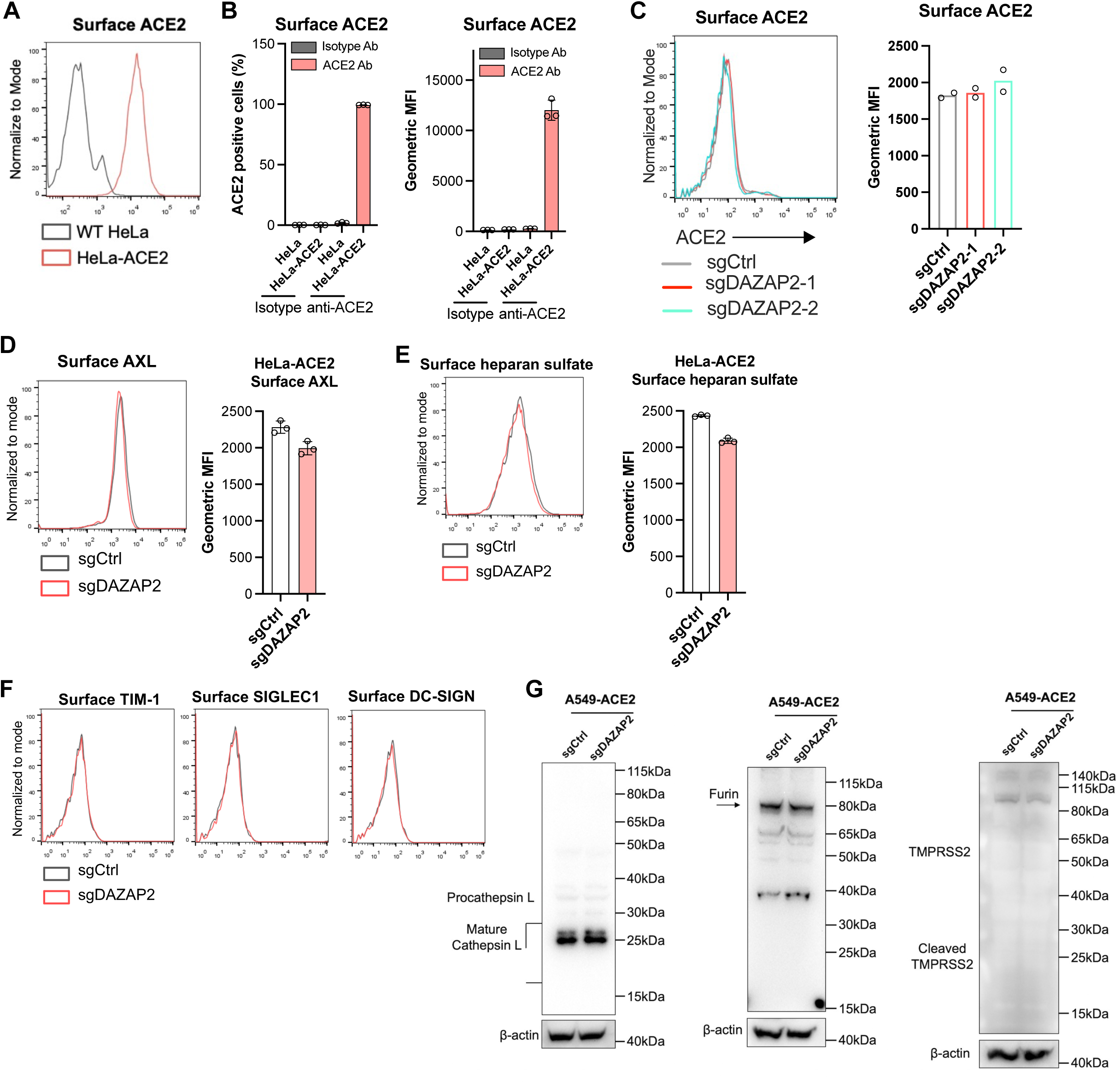
The expression of SARS-CoV-2 entry-related host factors. **A.** Flow cytometry analysis of surface expression of ACE2 in WT or ACE2-overexpressing HeLa cells stained with anti-ACE2 antibody. **B.** Flow cytometry analysis of surface expression of ACE2 in WT or ACE2-overexpressing HeLa cells stained with isotype or anti-ACE2 antibody. The percentage of ACE2 positive cells and geometric mean fluorescence intensity (MFI) were analyzed. **C-E.** Flow cytometry analysis of surface expression of ACE2, AXL, or heparan sulfate in A549-ACE2 (C) or HeLa-ACE2 (D and E) cells edited with control or *DAZAP2* sgRNA. The percentage of positive cells and geometric mean fluorescence intensity (MFI) were analyzed. **F.** Flow cytometry analysis of surface expression of TIM-1, SIGLEC1, or DC-SIGN in A549-ACE2 cells edited with control or *DAZAP2* sgRNA. **G.** Western blotting analysis of CTSL, Furin, or TMPRSS2 in A549-ACE2 cells edited with control or *DAZAP2* sgRNA.

## SUPPLEMENTARY TABLE LEGENDS

Supplementary Table 1. List of genes and scores after MaGeck analysis (see Excel file). Data was obtained by sequencing the sgRNAs from uninfected or sorted cells.

Supplementary Table 2. sgRNA sequences of genes selected for validation and other editing experiments (see Excel file).

Supplementary Table 3. List of primers and probes used for qRT-PCR experiments (see Excel file).

